# Novel and optimized mouse behavior enabled by fully autonomous HABITS: Home-cage Assisted Behavioral Innovation and Testing System

**DOI:** 10.1101/2024.09.29.615652

**Authors:** Bowen Yu, Penghai Li, Haoze Xu, Yueming Wang, Kedi Xu, Yaoyao Hao

**Affiliations:** The State Key Lab of Brain-Machine Intelligence, Zhejiang University, Hangzhou, China; Nanhu Brain-computer Interface Institute, Hangzhou, China; Department of Biomedical Engineering, Zhejiang University, Hangzhou, China; College of Computer Science and Technology, Zhejiang University, Hangzhou, China

**Keywords:** mouse cognition, autonomous training, behavioral innovation, machine teaching, behavioral optimization

## Abstract

Mice are among the most prevalent animal models used in neuroscience, benefiting from the extensive physiological, imaging and genetic tools available to study their brain. However, the development of novel and optimized behavioral paradigms for mice has been laborious and inconsistent, impeding the investigation of complex cognitions. Here, we present a home-cage assisted mouse behavioral innovation and testing system (HABITS), enabling free-moving mice to learn challenging cognitive behaviors in their home-cage without any human involvement. Supported by the general programming framework, we have not only replicated established paradigms in current neuroscience research but also developed novel paradigms previously unexplored in mice, resulting in more than 300 mice demonstrated in various cognition functions. Most significantly, HABITS incorporates a machine-teaching algorithm, which comprehensively optimized the presentation of stimuli and modalities for trials, leading to more efficient training and higher-quality behavioral outcomes. To our knowledge, this is the first instance where mouse behavior has been systematically optimized by an algorithmic approach. Altogether, our results open a new avenue for mouse behavioral innovation and optimization, which directly facilitates investigation of neural circuits for novel cognitions with mice.

## Introduction

Complex goal-directed behaviors are the macroscopic manifestation of high-dimensional neural activity, making animal training in well-defined tasks a cornerstone of neuroscience ^1–3^. Over the past decades, researchers have developed a diverse array of cognitive tasks and achieved significant insights into the underlying cognitive computations ^4^. Mouse has been increasingly utilized as model animal for investigating neural mechanisms of decision making, due to the abundant tools available in monitoring and manipulating individual neurons in intact brain ^5^. One notable example is the field of motor planning ^6^, which, after introduced into mouse model ^7^, was significantly advanced by obtaining causal results from whole brain circuits to genetics ^8^.

Traditionally, training mice in cognitive tasks was inseparable from human involvement in frequent handling and modifying shaping strategies according to the performance ^9^, thus labor-intensive and inconsistent as well as introducing unnecessary noise and stress ^10^. Recently, many works dedicated to design standard training setups and workflows, aiming for more stable and reproducible outcomes ^11–20^. For example, The International Brain Laboratory has recently showed mice can perform the task comparably across labs after a standard training within identical experimental setup ^19^. However, human intervention still has been a significant factor, which inevitable introduced the variability. Furthermore, training efficiency was still restricted by the limited experimental time as before, which necessitates motivational techniques like water restriction. The trainings were often restricted to a specific task and have not been extensively tested in other complex paradigms because of the requirement of prolonged training duration. These limitations highlighted the challenges to broadly and swiftly study comprehensive cognitive behaviors in mouse.

A promising solution to this scenario is to implement fully autonomous training systems. In recent years, researchers have focused on combining home-cage environments with automated behavioral training methodologies, offering a viable avenue to realize autonomous training ^21–34^. For instance, efforts have been made to incorporate voluntary head fixation within the home-cage to train mice on cognitive tasks ^21,27,29,30^. At the cost of increased training difficulty, they successfully integrate large-scale whole-brain calcium imaging ^21,29,30^ and optogenetic modulations ^27^ with fully automated home-cage behavior training. Moreover, other groups have conducted mouse behavior training in group-housed home-cages and utilizes RFID technology to distinguish individual mice ^28,29,32,34^. There are also studies which directly train freely moving animals in their home-cage to reduce stress and facilitate deployment ^10,25,28,31,32,34–36^. However, many of these studies have focused on single paradigms and incorporated complex components in their systems, which hindered high-throughput deployment for high-efficiency and long-term behavioral testing and exploring.

The training protocols employed in existing studies, no matter in manual or autonomous training, were often artificially designed ^19,27^, potentially failing to achieve optimal outcomes. For instance, a key issue in cognitive behavioral training is that mouse is likely to develop bias, i.e., obtaining reward only from one side. Various ‘anti-bias’ techniques ^19,27,37^ have been implemented to counteract the bias yet their efficacy in accelerating training or enhancing testing reliability remains unproven. From a machine learning standpoint, if we can accurately infer the animal’s internal models, it is possible to select a specific sequence of stimuli that will reduce the ‘teaching’ dimension of the animal and thus maximize the learning rate ^38^. Recently, an adaptive optimal training policy, known as AlignMax, was developed to generate an optimal sequence of stimuli to expedite the animal’s training process in simulation experiments ^39^. While many relative works have realized the theorical demonstration of the validity of machine teaching algorithms under specific conditions, these have been limited to teaching silicon-based learners in simulated environment ^39–41^. The direct application and empirical demonstration of these algorithms in real-world scenarios, particularly in teaching animals to master complex cognitive tasks, remain unexplored. There are two fundamental barriers for testing these algorithms in real animals training: firstly, traditional session-based behavioral training results in a discontinuous training process, introducing abrupt variation of learning rate and uncontrollable noise ^19,42^, which could undermine the algorithm’s capability; secondly, the high computation complexity of model fitting ^39,40^ poses challenges for deployment on microprocessor, thereby impeding extensive and high-throughput experiments. In the lack of human supervision, a fully autonomous behavioral training system necessitates an optimized training algorithm. Therefore, integrating fully automated training with machine teaching-based algorithms could yield mutually beneficial outcomes.

To address these challenges, we introduced a home-cage assisted behavioral innovation and testing system, referred as HABITS, which is a comprehensive platform featuring adaptable hardware and a universal programming framework. This system facilitates the creation of a wide array of mouse behavioral paradigms, regardless of complexity. With HABITS, we have not only replicated established paradigms commonly used in contemporary neuroscience research but have also pioneered novel paradigms that have never been explored in mice. Crucially, we have integrated a machine teaching-based optimization algorithm into HABITS, which significantly enhances the efficiency of both training and testing. Consequently, this study provides a holistic system and workflow for a variety of complex, long-term mouse behavioral experiments, which has the potential to greatly expand the behavioral reservoir for neuroscience and pathology research.

## Results

### System design of HABITS

The entire architecture of HABITS was comprised by two parts: a custom home-cage and behavioral components embedded in the home-cage **(Fig. 1A)**. The home-cage was made of acrylic plates with a dimension of 20×20×30 cm, which is more extensive than most of commercial counterparts for single-housed mouse. A tray was located at the bottom of the cage where *ad libitum* food, bedding, nesting material (cotton) and enrichment (a tube) were placed **(Fig. 1B)**. Experimenters can change the bedding effortlessly just by exchanging the tray. The home-cage also included an elevated, arc-shaped weighting platform inside, providing a lose body constraint for the mouse during task performing (**Fig. 1A)**. Notably, a micro load cell was installed beneath the platform, which can readout body weight of the mouse for long-term health monitoring. The cage was compatible with the standard mouse rack and occupied as small space as standard mouse cage **(Fig. 1C)**.

**Fig. 1.**
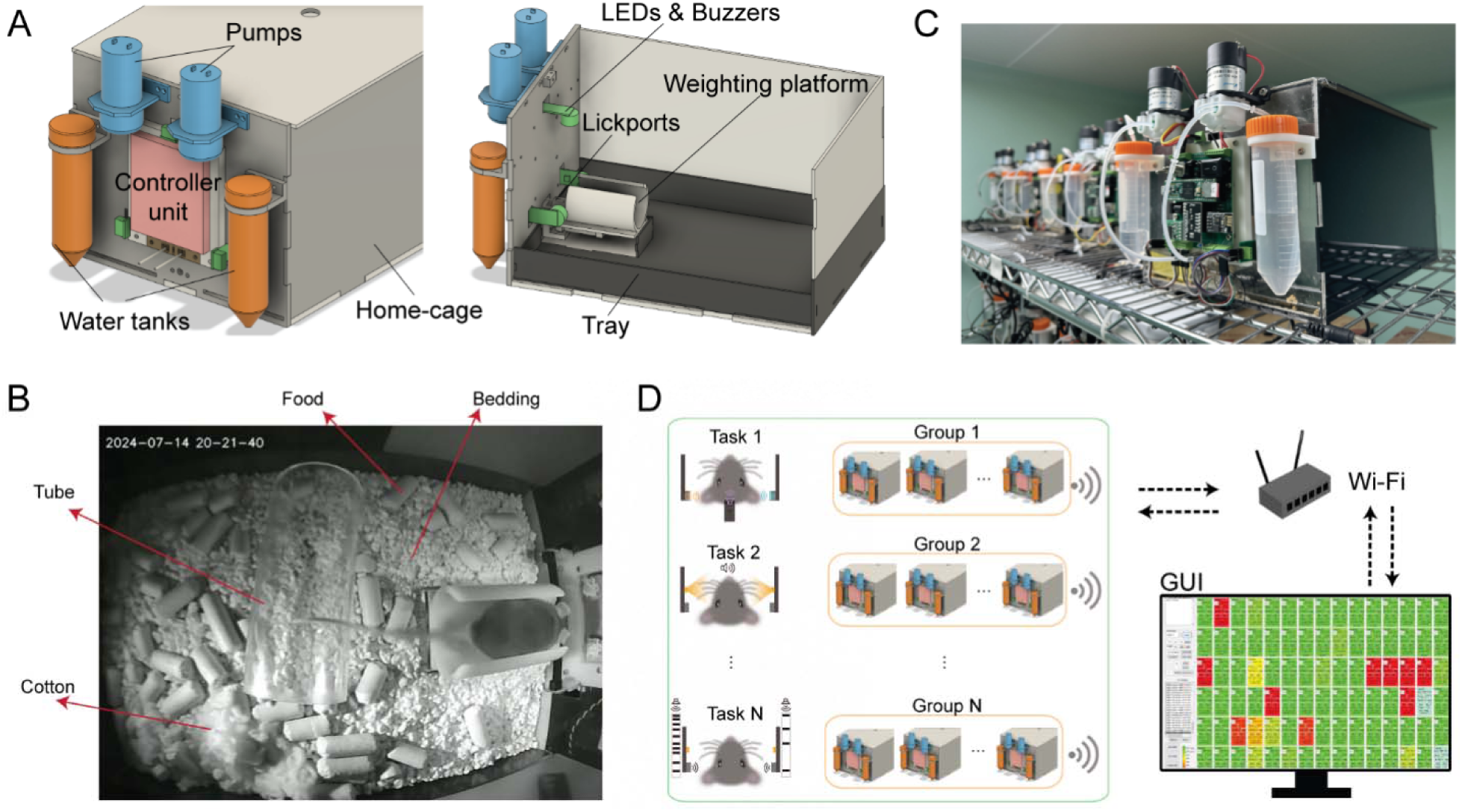
System setup for HABITS. (**A**), Front (left) and side (right) view of HABITS, showing components for stimulus presenting (LEDs & buzzers), rewarding (water tanks and pumps), behavioral reporting (lickports) and health monitoring (weight platform). These components are coordinated by controller unit and integrated into mouse home-cage with a tray for bedding change. (**B**), HABITS installed on standard mouse cage rack. (**C**), Mouse, living in home-cage with food, bedding, nesting material (cotton) and enrichment (tube), is performing task on the weight platform. (**D**), System architecture for high-throughput behavioral training, showing different tasks are running in parallel groups of HABITS, which further wirelessly connect to one single PC through Wi-Fi to stream real-time data to the graphic user interface (GUI).

To perform behavioral training and testing in HABITS, we constructed a group of training-related components embedded in the front panel of the home-cage **(Fig. 1A)**. Firstly, three stimulus modules (LEDs and buzzers) for light and sound presenting were protruded from front panel and placed in the left, right and top position around the weighting platform, enabling generation of visual and auditory stimulus modalities in three different spatial locations. The mouse reported the decisions about the stimuli by licking either left, right or middle lickports installed in the front of the weighting platform. Finally, peristaltic pumps draw water from water tanks into lickports, serving as the reward for the task, which was the sole water source for the mouse throughout the period living in the home-cage. In the most common scenario, mouse living in home-cage stepped on the weighting platform and triggered trials by licking on the lickports to obtain water **(Fig. 1B, Movie S1)**.

To endow autonomy to HABITS, a microcontroller was used to interface with all training-related components and coordinate the training procedure **(Fig. S1A)**. We implemented a microcontroller-based general programming framework to run finite state machine with millisecond-level precision (see Methods). By using the APIs provided by the framework, we can easily construct arbitrarily complex behavioral paradigms and deploy them into HABITS **(Fig. S1B)**. Meanwhile, the paradigms were usually divided into small steps from easy to hard and advanced according to the performance of the mouse. Another important role of the microcontroller was to connect the HABITS to PC wirelessly and stream real-time behavioral data via a Wi-Fi module. A graphic user interface (GUI), designed to remotely monitor each individual mouse’s performance, displayed history performance, real-time trial outcomes, body weights, etc. **(Fig. S1C)**. Meanwhile, the program running in the microcontroller could be updated remotely by the GUI when changing mouse and/or task. As a backup, information of training setup, task parameters and all behavioral data were also stored in a SD card for offline analysis.

To increase the throughput of behavioral testing, we built more than a hundred of independent HABITS and installed them on standard mouse racks **(Fig. S1D)**. The total material cost for each HABITS was less than $100 **(Table SI)**. All HABITS, operating different behavioral tasks across different cohorts of mice, were organized into groups according to the tasks, and wireless connected to a single PC **(Fig. 1D)**. The states of each individual HABITS can be accessed and controlled remotely by monitoring the corresponded panels in GUI, thereby significantly improving the experimental efficiency. We developed a workflow to run behavioral training and testing experiment in HABITS **(Fig. S1E)**. Firstly, initiate HABITS system by preparing the home-cage, deploying training protocols and weighting the naive mouse that did not need to undergo any water restriction. Then, mouse interacted with fully autonomous HABITS at their own pace without any human intervention. In this study, mice housed in HABITS went through 24/7 behavioral training and testing for up to 3 months **(Movie S2)**, though longer duration was supported. Finally, data were harvested from SD card and offline analyzed.

Therefore, HABITS permitted high-throughput, parallel behavioral training and testing in a fully autonomous way, which, contrasting with manual training, allows for possible behavioral innovation and underlining neural mechanism investigation.

### HABITS performance probed by multimodal d2AFC task

We next deployed a well-established mouse behavioral paradigm, delayed two-alternative forced choice (d2AFC), which was used to study motor planning in mouse ^7,43^, in HABITS to demonstrate the performance of our system **(Fig. 2).**

**Fig. 2.**
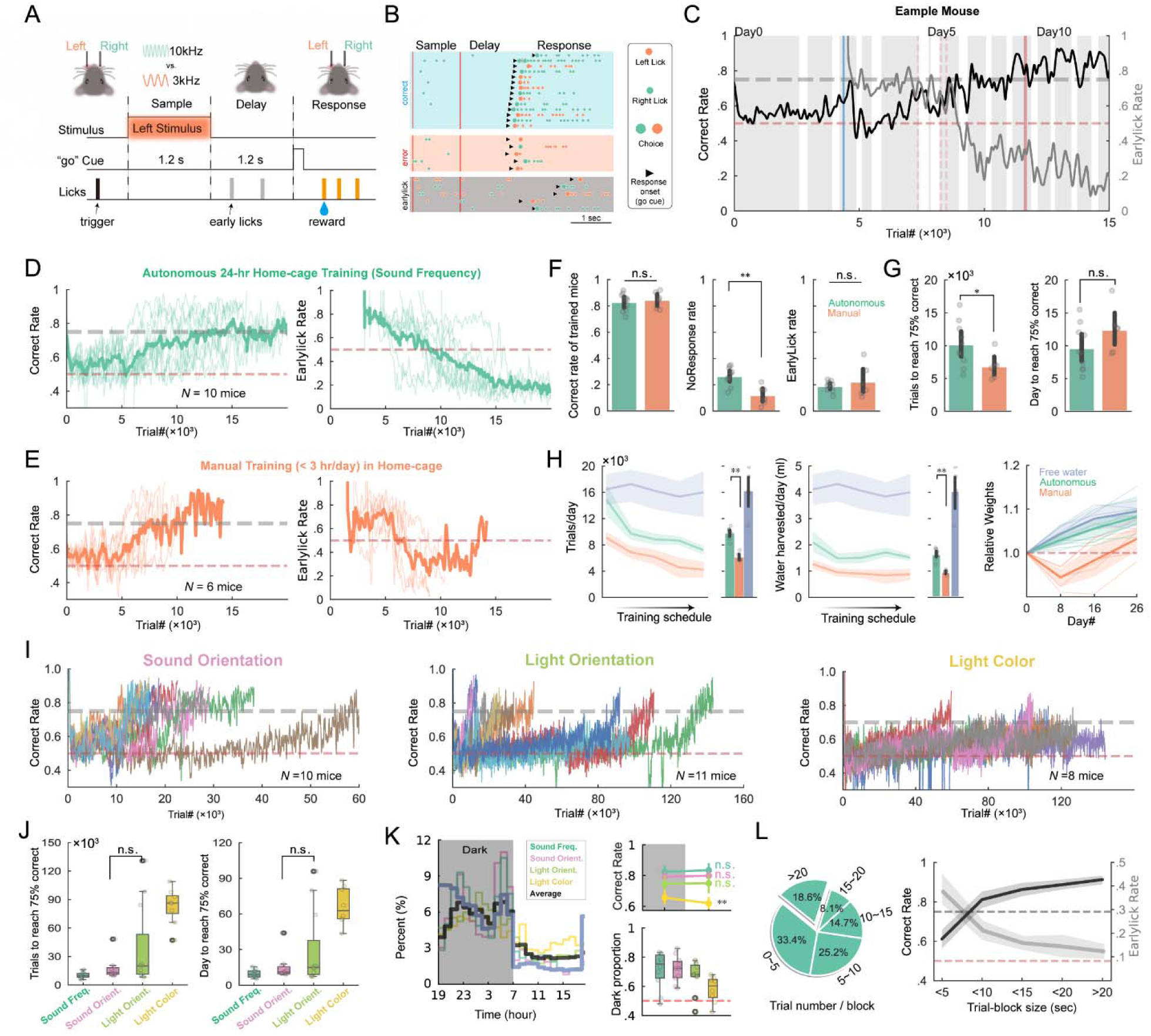
HABITS performance in d2AFC task. **(A)**, Task structure for d2AFC based on sound frequency. **(B)**, Example licks for correct (blue), error (red) and earlylick (gray) trials. Choice is the first lick after response onset. **(C)**, Correct rate (black line) and earlylick rate (grey line) of an example mouse during training in HABITS for the first 13 days. Shaded blocks indicate trials occurred in dark cycle. Trials with earlylick inhabitation only occur after blue vertical line. Red vertical dash lines represent delay duration advancement from 0.2 s to 1.2 s. **(D)**, Averaged correct rate (left) and earlylick rate (right) for all mice trained in d2AFC. Criterion level (75%) and chance level (50%) are labeled as gray and red dash lines, respectively. **(E)**, Same as (D) but for manual training (1∼3 hour/day in home-cage). **(F)**, Averaged correct rate (left), earlylick rate (middle) and no response rate (right) of expert mice trained with the two protocols. **(G)**, Averaged number of trials (left) and days (right) to reach the criterion performance for the two training protocols. Circles, individual mice. Errorbar, mean and 95% CI across mice. **(H)**, Left, number of trials performed per day throughout the training schedule for three different protocols. Error bar indicates the mean and 95% confidence interval (CI) across mice. Middle, volume of water harvested per day. Right, Relative body weights of mice in day 0, 8, 16, 26. Bold line and shades indicate mean and 95% CI across mice. **(I)**, Behavioral performance of all mice training in d2AFC task based on sound orientation (left), light orientation (middle), and light color (right). **(J)**, Box plot of average number of trials (left) and days (right) to reach the criterion performance for d2AFC tasks with different sensory modalities. **(K)**, Left, percentage of trials performed as a function of time in a day for the four modalities trained autonomously (thick black shows the average). Shaded area indicates the dark cycle. Top right, averaged correct rate of grouped mice in dark cycle versus light cycle. Error bars show 95% CI across mice. Bottom right, box plot of the averaged proportion of trials performed in dark cycle for the four modalities. Data collected from expert mice. **(L)**, Left, percentage of trials in blocks with varying number of consecutive trails for automated training in home-cage. Right, correct rate and earlylick rate as functions of trial block size. Grey dash line, the criterion performance; Red dash line, chance performance level. Data collected from trials of expert mice. For significance levels not mentioned in all figures, n.s., not significant, p> 0.05; *, p<0.05; **, p<0.01 (two-sided Wilcoxon rank-sum tests).

Mice, living in HABITS all the time, licked any of the lickports to trigger a trial block at any time. In d2AFC task with sound frequency modality **(Fig. 2A)**, mice needed to discriminate two tone frequency presented at the beginning of the trial for a fixed duration (sample epoch, 1.2 sec) and responded the choice by licking left (low tone) or right (high tone) lickports following a brief ‘go’ cue after a fixed delay duration (1.2 sec). Licks during delay epoch were prohibited and unwished lick (early licks) will pause the trial for a while. Correct choices were rewarded, while incorrect choices resulted in noise and timeout. If mice did not lick any of the lickports after ‘go’ cue for a fixed period (i.e., no-response trial), the trial block was terminated. The next trial block was immediately entering the state of to be triggered. Mice can only learn the stimulus-action contingency by trial-and-error. To promote learning, we designed an algorithm comprised of many subprotocols to shape the behavior step by step **(Fig. S2A)**. **Fig. 2B** illustrated example licks during correct, error and early lick trials for the task. As training progressed, the correct rate increased and earlylick rate declined gradually for the example mouse within the first 2-week **(Fig. 2C)**. All the 10 mice enrolled in this task effectively learned the task and suppressed licking during the delay period within 15 days **(Fig. 2D)**, achieving an average of 980 ± 25 (mean ± SEM) trials per day.

We also tested another training scheme, i.e., limited duration per day in HABITS, to simulate the situation of manual training with human supervision. These mice were water restricted and manually transferred from traditional home-cage to HABITS daily for 1-3 hours training session. The duration of sessions was determined by the speed of harvesting water as we controlled the daily available volumes of water to approximately 1 ml ^9^. All the 6 mice learned the task as the autonomous counterpart **(Fig. 2E)** with similar final correct and earlylick rate (except that the no-response rate was significantly lower for manual training) **(Fig. 2F)**. Logistic regression of the mice’s choice revealed similar behavioral strategies were utilized throughout the training for both group (**Fig. S2B-D**). Autonomous training needed significantly more trials than manual training (10,164 ± 1,062 vs. 6,845 ± 782) to reach the criterion performance, however, the number of days was slightly less due to the high trial number per day (**Fig. 2G**). As shown in **Fig. 2H**, autonomous training yielded significantly higher number of trial/day (980 ± 25 vs. 611 ± 26, Fig. 2H left) and more volume of water consumption/day (1.65 ± 0.06 vs. 0.97 ± 0.03 ml, Fig. 2H middle), which resulted in monotonic increase of body weight that was even comparable to the free water group (Fig.2H right). In contrast, the body weight in manual training group experienced a sharp drop at the beginning of training and was constantly lower than autonomous group throughout the training stage (**Fig. 2H right**). As the training advanced, the number of trials triggered by mice per day decreased gradually for both groups of mice, but the water consumption per day kept relatively stable. At the end of manual training, we transferred all mice to autonomous testing, and found that the number of trial and consumption water per day dramatically increased to the level of the autonomous training, suggesting mice actually needed more water throughout the day (**Fig. S2E**). These results indicated that autonomous training achieved similar training performance as manual training and maintained a more health state of mice.

Three more cohorts of mice were used in d2AFC tasks with different modalities, which included sound orientation (left vs. right sound), light orientation (left vs. right light) and light color (red vs. blue). Using the same training protocol as in the sound frequency modality, we trained 10, 11 and 8 mice on these tasks, respectively **(Fig. 2I)**. Mice required different amounts of trials or days to discriminate these modalities, with light color discrimination being the most challenging (an average of 82,932 ± 6,817 trials), consistent with the limited sensitivity to light wavelength of mice (**Fig. 2J**). We also tried other modalities, like blue vs. green and flashed blue vs. green, but all failed (see **Table I**).

**Table 1.** All tasks training in HABITS.

| Protocol name (abbreviation) | Modality | Animals trained (trained / used) | Note |
| --- | --- | --- | --- |
| delayed 2-Alternative Forced Choice (d2AFC) | Sound frequency (3k vs. 10 kHz) | 11/11 | Figure 2 |
|  | Sound orientation (Left vs. Right) | 10/11 |  |
|  | Light orientation (Left vs. Right) | 10/11 |  |
|  | Light color (Blue VS. Red) | 8/11 |  |
|  | Light color (Green VS. Blue) | 0/10 | Mice are insensitive to light colors. |
|  | Light color (flashed Green VS. Blue) | 0/10 |  |
| Reaction time 2AFC (RT-2AFC) | Sound frequency (3k vs. 12kHz) | 6/6 | Figure S3 |
| Contingency reversal | RT-2AFC, sound frequency (3k vs. 12kHz) | 8/8 | Figure 3A |
| Continuous learning | Sound freq. (3k vs. 12kHz), reversal sound freq., sound orient. (Left vs. Right), reversal sound orient., light orient. (Left vs. Right) | 10 /30 | Figure 4A;<br>20 mice did not learn light oriental modal within 90 days. |
| Evidence accumulation with spatial cue | Poisson distributed clicks with spatial diff. (Left vs. Right) | 8/10 | Figure 3C |
| Multimodal Integration | Poisson distributed clicks and flashes (4 vs. 20 events/s) | 13/15 | Figure 3D |
|  | Poisson distributed light flashes (4 vs. 20 events/s) | 3/15 | 12/15 mice failed to discriminate light flash |
| Confidence probing task | Sound frequency (8k vs. 32kHz) | 5/6 | Figure 3E |
|  | Sound frequency (8k vs. 32kHz) and Poisson distributed clicks with spatial diff. (Left vs. Right) | 0/12 | Delay period up to 8 sec, failed |
| Value-based dynamic foraging task | No sensory cues (Block size from 500 to 100) | 6/6 | Figure S4 |
| Working memory task | Temporal regular clicks with 5 alternative rates (8, 16, 32, 64, 128Hz) | 5/6 | Figure 3B |
|  | Temporal regular clicks with 3 alternative rates (8, 32, 128Hz) | 8/8 | Figure S5 C |
| double Delayed Match Sample (dDMS) | Sound frequency (3k & 12kHz) | 10/10 | Figure 4B;<br>Sample & test period: 500ms |
|  | Sound frequency (3k & 12kHz) | 0/10 | Random sample & test period: 100 or 1000ms |
| delayed 3-Alternative Forced Choice (d3AFC) | Sound frequency (8k vs. 16k vs. 32kHz) | 14/14 | Figure 4C |
|  | Sound frequency reversal (8k vs. 16k vs. 32kHz) | 4/4 | Figure S5B |
|  | Sound orientation (Left vs. Middle vs. Right) | 6/6 | Figure S5A |
| Context-dependent attention task | Sound frequency (3k vs. 12kHz) or sound orientation (Left vs. Right); regular click rates (16 vs. 64Hz) as context cue | 6/6 | Figure 4D |
|  | Sound orientation (Left vs. Right) or light orientation (Left vs. Right); regular click rates (16 vs. 64Hz) as context cue | 0/10 | Mice failed in light modality |
|  | Sound orientation (Left vs. Right) or light orientation (Left vs. Right); Sound frequency (3k vs. 12kHz) as context cue | 0/20 |  |
| Machine teaching algorithm | Working memory task, Temporal regular clicks with 5 alternative rates and full stimulus matrix (8, 16, 32, 64, 128Hz) | 7/7 (MT)<br>5/8(Random) | Figure 5B |
|  | RT-2AFC, Sound frequency (3k vs. 12kHz) | 10/10 (MT)<br>10/10 (random)<br>10/10 (antibias) | Figure 5F;<br>Same group of mice used in continuous learning |
|  | 2AFC with Sound frequency (3k vs. 12kHz), sound orientation (Left vs. Right) and sound orientation reversal, respectively | 10/10 (MT)<br>8/8 (random) | Figure 6 |
| **Total** | N/A | **200/284** | N/A |

The learning rate between sound and light orientation discrimination tasks was similar (p = 0.439, two-sided Wilcoxon rank-sum test), but the variation for light orientation was large, indicating possible individual differences (**Fig. 2J**). All modalities maintained good health state indicated by the body weight for up to 2 months (**Fig. S2F**). Another two types of 2AFC, reaction time ^44^ (RT-2AFC, **Fig. S3**) and random foraging (**Fig. S4**) task, were also successfully tested in HABITS. Notably, the dynamic foraging task ^45,46^, which heavily relies on historical information, was first demonstrated in a fully autonomous training scheme for freely moving mice with similar block size and performance.

Two important behavioral characteristics were revealed across all the modalities in 2AFC task. Firstly, as our high-throughput behavioral training platform operated on a 12:12 light-dark cycle, the long-term circadian rhythm of mice can be evaluated based on the number of triggered trials and performance during both cycles. We found all mice exhibited clear nocturnal behavior with peaks in trial proportion at the beginning and end of the dark period (**Fig. 2K**), which was consistent with previous studies ^25,27^. The light color modality exhibited slight lower percentage of trials during dark (57.89% ± 3.18% vs. above 66% for other modalities) (**Fig. 2K**), possibly the light stimulus during trials affected the circadian rhythms of the mice. It was also observed that all mice except the light color modality showed no significant differences in correct rate between light and dark cycle after they learned the task (**Fig. 2K**). The higher performance in dark environment for light color modality implied that light stimulus presented in dark environment was with higher contrast and thus better discernibility. Secondly, as we organized the trials into blocks, the training temporal dynamic at trial-level could be examined. We found more than two-third of the trials was done in >5-trial blocks (**Fig. 2L left**) which resulted in more than 55% of the trials were with inter-trial-interval less than 2 seconds (Fig. S2H). The averaged block duration was 27.64 ± 1.73 sec and mice triggered another block of trials within 60 sec in more than 60% of cases. Meanwhile, we also found that the averaged correct rate increased and the earlylick rate decreased as the length of block increased (**Fig. 2L right**), which suggested that mice were more engaged in the task during longer blocks.

These results showed that mice can learn and perform cognitive tasks in HABITS with various modalities in a fully autonomous way. During the training process, mice maintained good health condition, although without any human intervention. Due to the high-efficiency training with less labor and time, it gave us an opportunity to explore and study more widespread cognitive behavioral tasks in mice.

### Various cognitive tasks demonstrated in HABITS

We next tested several representative tasks that were commonly used in the field of cognitive neuroscience to demonstrate the universality of HABITS. It is worth noting that many new features of the behavior could be explored due to the autonomy and advantages of HABITS, in terms of either quantity or quality **(Fig. 3)**.

**Fig. 3.**
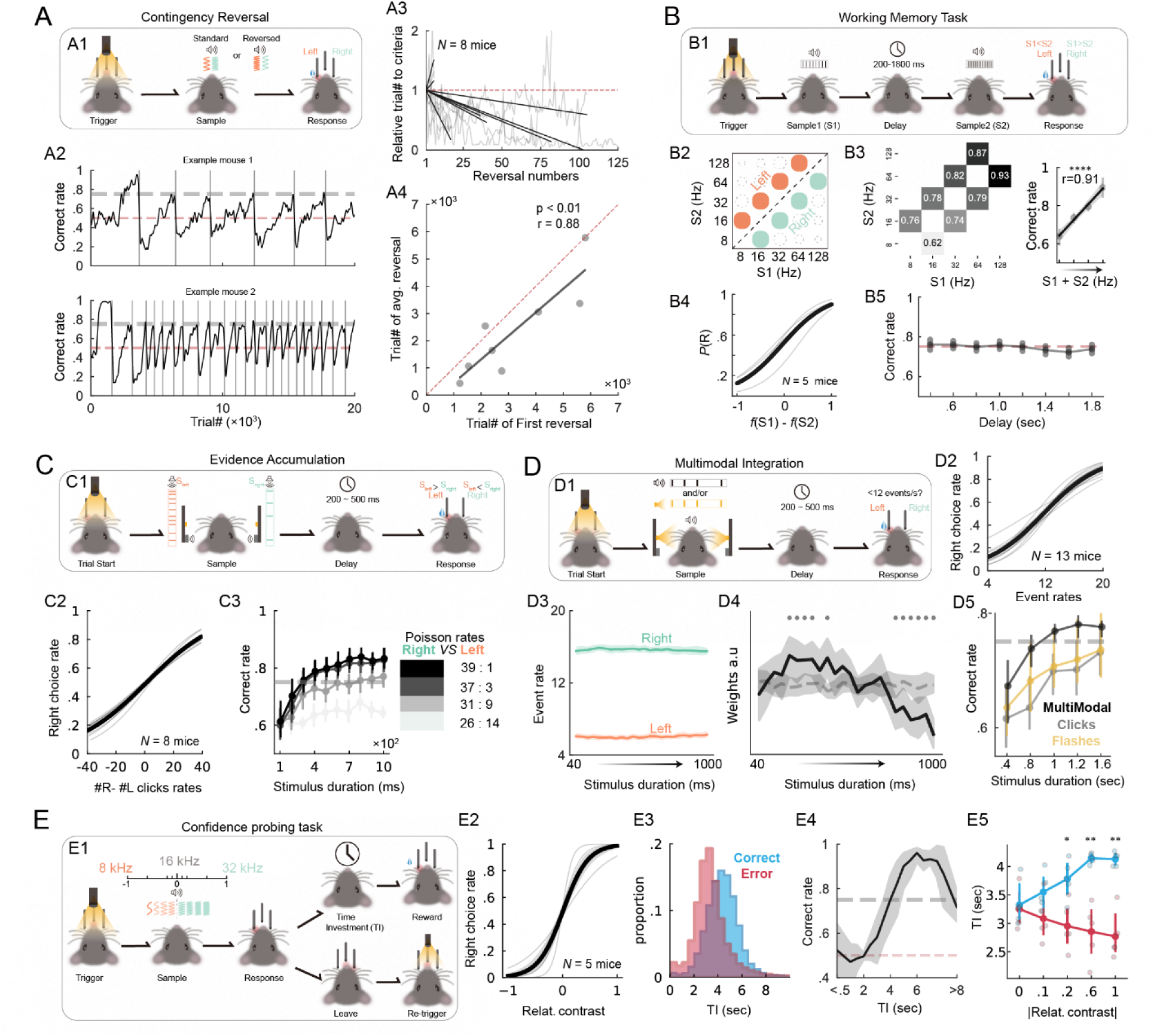
Representative cognitive task performed in HABITS. (**A**), Contingency reversal task. (*A1*), Task structure. (*A2*), Correct rate of example mice with different learning rates. Grey vertical lines indicate contingency reversal. (*A3*), Relative number of trials to reach the criterion as a function of reverse times. Grey lines, individual mice. Black lines, linear fit. (*A4*), Number of trials in the first reversal learning versus the average number of trials of the rest of contingency reversal learning for each mouse (each dot). Black line, linear regression. Red dash line, diagonal line. (**B**), Working memory task with sound frequency modality. (*B1*), Task structure. (*B2*), Stimulus generation matrix (SGM) for left (orange) and right (green) trials. (*B3*), left, averaged correct rate for each stimulus combination tested. Right, averaged correct rate for each (S1+S2) stimulus combination across mice. Black line and shade, linear regression and 95% CI. (*B4*), Averaged psychometric curves, i.e., percentage of right choice as a function of frequency difference between sample1 and sample2. (*B5*), Averaged correct rate as a function of delay duration. **(C)**, Evidence accumulation with spatial cue task. (*C1*), Task structure. (*C2*), Averaged psychometric curves, i.e., performance as a function of the difference between right and left clicks rates. (*C3*), Averaged correct rate across all mice as a function of sample duration for different Poisson rates (different colors). Error bar represents 95% CI. **(D)**, Multimodal integration task. (*D1*), Task structure. (*D2*), Averaged correct rate across all mice as a function of sample duration for different stimulus modalities (different colors). (*D3*), Averaged event rates during sample period for left (red) and right (blue) choice trials. (*D4*), Averaged weights (black line) of logistic regression fitting to the choice of trials across expert mice tested in > 1000 trials (N = 11 mice) from the first bin (40ms) to the last bin (1000ms) of the sample period. Gray dash line represents null hypothesis. Grey dots indicate significance, p<0.05, two-sided *t*-tests. (*D5*), Psychometric curve for trials with multimodal stimulus. **(E),** Confidence probing task. (*E1*), Task structure. (*E2*), Psychometric curve, i.e., right choice rate as a function of relative contrast (log scaled relative frequency). (*E3*), Histogram of time invested (TI) for both correct and error trials. (*E4*), Averaged correct rate across all mice as a function of TI. (*E5*), Averaged TI as a function of absolute relative contrast for both correct and error trials. Cycles, individual mice; *, p<0.05; **, p<0.01, two-sided Wilcoxon rank-sum tests.

Contingency reversal task was cognition-demanding task and previously used to investigate cognitive flexibility ^47^ **(Fig. 3A)**. In the task, the contingency for reward switched without any cues once mice hit the criteria performance (**Fig. 3A1**). Mice can dynamically reverse their stimulus-action contingency, though with different learning rate across individuals **(Fig. 3A2)**. Given the advantages of long-term autonomous training in HABITS, all mice undergone contingency reversal for an average of 52.25±16.39 times and one mouse achieved up to 125 times in 113 days. Most of mice (6/8) gradually decreased the number of trials to reach criterion across multiply contingency reversal representing an effect of learning to learn ^45,48^ **(Fig. 3A3)**. Meanwhile, the average number of trials to reach criterion during the reversal was highly correlated with the trial number in the first reversal learning which represented the initial cognitive ability or the sensory sensitivity of individual mice **(Fig. 3A4)**.

Working memory was another important cognition that were vastly investigated using rodent model with auditory and somatosensory modalities ^48,49^. Here we utilized a novel modality, regular sound clicks, to implement a self-initiated working memory task, which required mice to compare two click rates separated by a random delay period **(Fig. 3B1)**. We initially validated that mice did employ a comparison strategy in a 3×3 stimulus generation matrix (SGM), instead just taking the first cue as a context **(Fig. S5A)**. Subsequently, we expanded the paradigm’s perception dynamics to 5×5, and reduced the relative perceptual contrast between two neighboring stimuli to one octave **(Fig. 3B2)**. HABITS enabled investigation of detailed behavioral parameters swiftly. We noticed that the discrimination ability for mice was significantly better in higher frequency range **(Fig. 3B3)**, which may be caused by the different sensitivity of mice across the spectrum of regular click rate. During testing stage, we varied the contrast of the preceding stimulus while maintaining the succeeding one; the psychometric curves affirmed mice’s decision-making based on perceptual comparison of two stimuli and validated their perceptual and memory capacities **(Fig. 3B4)**. Meanwhile, mice could maintain stable working memory during up to 1.8 sec delay, demonstrating mice can perform this task robustly **(Fig. 3B5)**.

Evidence accumulation introduced more dynamics to the decision-making within individual trials and were widely utilized ^50–52^. In the task implemented in HABITS, mice needed to compare the rate of sound clicks randomly appeared over two sides during sample epoch and made decision following a ‘go’ cue after a brief delay **(Fig. 3C1)**. We successfully trained 8/10 mice to complete this task as revealed by the psychometric curves **(Fig. 3C2)**. After mice learned the task, we systematically tested the effect of evidence accumulation versus the task performance. All mice exhibited consistent behavioral patterns which was correlated with the evidence of stimulus, i.e., longer sample period and/or higher stimulus contrast led to higher performance **(Fig. 3C3)**. It showed an evident positive correlation between evidence accumulation and task performance.

Multimodal integration ^53–55^ was also tested in HABITS with sound clicks and light flashes as dual-modality events in the evidence accumulation framework. Mice were required to differentiate whether event rates were larger or smaller than 12 event/sec **(Fig. 3D1)**. We successfully trained 13/15 mice in this paradigm with multimodal or sound clicks stimulus **(Fig. 3D2)**, and only 3 mice achieved performance criteria in trials with light flashes stimulus. Since the events were presented non-uniformly within each trial, we wondered the dynamics of decision-making process along the trial. Firstly, we divided all trials with multimodal stimulus into two groups according to the choice of expert mice. The uniform distribution of events within a trial indicated that mice considered the whole sample period to make decision **(Fig. 3D3)**. Secondly, we used a logistic regression model to illustrate the mice’s dependency on perceptual decisions throughout the entire sample period. We found that mice indeed tended to favor earlier stimuli (i.e., higher weights) in making their final choices **(Fig. 3D4)**, consistent with previous research findings ^54^. Lastly, we further tested modulated unimodal stimuli with different sample period in testing stage, in which accuracy correlated positively with sample period and conditioned test for different combination of modalities demonstrated evidence accumulation and multimodal integration effect, respectively. **(Fig. 3D5)**.

Confidence was another important cognition along with the process of decision-making, which was investigated in rat ^56,57^ and more recently in mice ^58^ model previously. We introduced confidence-probing task in HABITS **(Fig. 3E)**, in which mice needed to lick twice the correct side for acquiring a reward; the two licks were separated by a random delay during which licking at other lickports prematurely ended the trial **(Fig. 3E1)**. This unique design connects mice’s confidence about the choice, which was hidden, with the time investment (TI) of mice between two licks, which was an explicit and quantitative metric. We successfully trained 5/6 mice **(Fig. 3E2)**, and most importantly, there was a noticeable difference of TI in correct versus incorrect trials **(Fig. 3E3)**. In detail, trials with longer TIs tended to have higher accuracies **(Fig. 3E4)**. Furthermore, the TI was also modulated by the contrasts of stimuli; as contrast decreased, mice exhibited reduced confidence about their choices, manifesting as decreased willingness to wait in correct trials, and conversely in error trials **(Fig. 3E5)**.

In summary, mice could undergo stable and effective long-term training in HABITS with various cognitive task commonly used in state-of-the-art neuroscience. These tasks running in HABITS were demonstrated to exhibit similar behavioral characteristics to previous studies. In addition, some new aspects of the behavior could be systematically tested in HABITS due to its key advantage of autonomy. This high level of versatility, combined with the ability to support arbitrary paradigm designs, suggests that more specialized behavioral paradigms could potentially benefit from HABITS to enhance experimental novelty.

### Innovating mouse behaviors in HABITS

One of the main goals of HABITS was to expand mouse behavioral reservoir by developing complex and innovative paradigms that had previously proven challenging or even impossible for mice. These paradigms imposed higher cognitive abilities demands, which required extensively long period to test in mouse model. HABITS enabled unsupervised, efficient and standardized training of these challenging paradigms at scale, and thus was suitable for behavioral innovations.

Firstly, leveraging the autonomy of HABITS, we tested mouse’s ability to successively learn up to 5 tasks one after another without any cues **(Fig. 4A)**. These tasks included 2AFC based on sound frequency, sound frequency reversal, sound orientation (pro), sound orientation (anti), and light orientation (**Fig. 4A1**). Firstly, the results showed that mice could quickly switch from one task to another and the learning rates across these sub-tasks roughly followed the learning difficulty of modalities **(Fig. 4A2)**. Specially, reversal of sound frequency was cognitively different from reversal of sound orientation (i.e., from pro to anti) which resulted in significant longer learning duration (**Fig. 4A2**). Secondly, mice dealt with new tasks with higher reaction time and gradually decreased as training progressed **(Fig. 4A3)**. It implied a uniform strategy mice applied: mice chose to respond more slowly in order to learn quickly ^59^. Lastly, mice exhibited large bias at the beginning of each task in all task including task without reversals **(Fig. 4A4)**. This means that mice acquired reward only from one lickport in the early training and switched strategy to follow current stimulus gradually, which implied a changing strategy from exploration to exploitation.

**Fig. 4.**
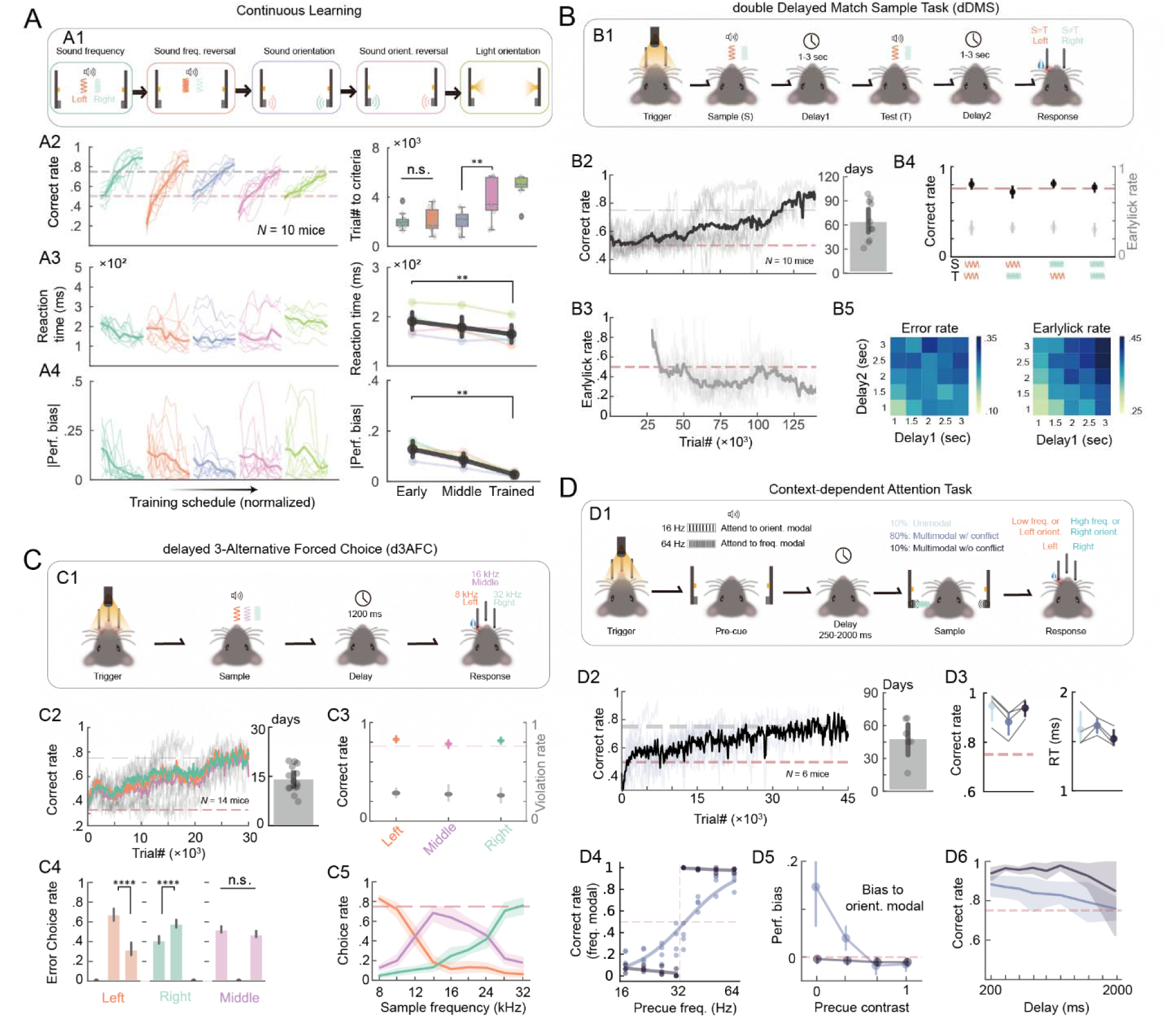
Challenging mouse tasks innovated in HABITS. (**A**), Continuous learning task. (*A1*), Task structure showing mice learning five subtasks one by one. (*A2*), Left, averaged correct rate of all mice performing the five tasks (different colors) continually. All task schedules are normalized to their maximum number of trials and divided to 10 stages equally. Right, box plot of number of trials to criteria for each task. (*A3*), Left, averaged reaction time of all mice performing the five tasks continually. Right, averaged median reaction time across the five tasks during early (perf. < 0.55), middle (perf. < 0.75) and trained (perf. > 0.75) stage. Error bar indicates 95% CI. (*A4*), Same as (A3) but for absolute performance bias. n.s., p>0.05; **, p<0.01, two-sided Wilcoxon signed-rank tests. **(B),** Double delayed match sample task (dDMS) with sound frequency modality. (*B1*), Task structure. (*B2*), Averaged correct rate across all mice during training (left) and averaged number of days to reach the criterion (right). (*B3*), Averaged earlylick rate across all mice. (*B4*), Averaged correct rate (black) and earlylick rate (gray) for all combination of sample and test stimulus. (*B5*), Heatmap of error rate (left) and earlylick rate (right) varies with different combination of delay1 and delay2 durations. **(C),** Delayed 3 alternative forced choice (d3AFC). (*C1*), Task structure. (*C2*), Averaged correct rate across all mice during training (left, colors indicate trial types) and averaged number of days to reach the criterion performance (right). (*C3*), Averaged correct rate (colors indicate trial types) and earlylick rate (gray) for different trial types. (*C4*), Averaged error rate of choices conditioning trial types. In each subplot, the position of bars corresponds to different choices. ****, p<0.0001, n.s., p>0.05, two-sided t-tests. (*C5*), Averaged choice rates for the three lickports (colors) as a function of sample frequency. Data collected from trained mice. **(D),** Context-dependent attention task. (*D1*), Task structure. (*D2*), Averaged correct rate across all mice during training (left, data only from trials with multimodal w/ conflict) and averaged number of days to reach the criterion (right). (*D3*), Correct rate (left) and reaction time (right) conditioning modalities. (*D4*), Averaged psychometric curve and partitioned linear regression for the multimodal with and without conflict conditions, respectively. (*D5*), Performance bias to sound orientation modal as a function of pre-cue contrast, for the two multimodal conditions. (*D6*), Averaged correct rate as a function of delay duration.

Delayed match sample task was quite challenging for mice, and only olfactory and tactile modalities were implemented previously ^60–62^. Recently, auditory modality was introduced but only in go/no-go paradigm ^63^. We next constructed a novel double delayed match sample task (dDMS) task (**Fig. 4B**), which required mice to keep working memory of first sound frequency (low or high) during the first delay, match to the second sound based on XOR rules, make a motor planning during the second delay, and finally make a 2AFC choice **(Fig. 4B1)**. All the 10 mice achieved the performance criteria during the automated training process, though, an averaged 64.45 ± 7.88 days was required, which was equivalent to more than 120,000 trials (**Fig. 4B2-3**). After training, the four trial types (i.e., four combinations of frequencies) achieved equally well performance **(Fig. 4B4)**. During testing stage, we systematically randomized the duration of the two delays (ranging from 1 to 3 sec), and revealed increased error rates and earlylick rates as the delay increased **(Fig. 4B5)**. Challenging tasks which demanded months of training were well suitable for HABITS, otherwise, difficult or even impossible for manual training.

Subsequently, we attempted to expand the choice repertoire of d2AFC into 3-alternative forced choice (d3AFC), utilizing the three lickports installed in HABITS **(Fig. 4C)**. Previous studies have implemented multiple choice tasks, but only based on spatial modalities ^64–66^. In our system, mice learned to discriminate low (8 kHz), medium (16 kHz) and high (32 kHz) sound frequencies and lick respectively left, middle and right lickport to get reward **(Fig. 4C1)**. Mice needed to construct two psychological thresholds to conduct correct choices. We successfully trained all the 14 mice to perform the tasks in 13.85 ± 1.05 days, with similar performance among the left, middle and right trial types **(Fig. 4C2)**. The final correct rate and early lick rate made no differences for the three trial types (**Fig. 4C3**). Interestingly, mice made error choices more in the most proximity lickport; for the middle trials, mice made error choice equally in the left and right side (**Fig. 4C4**). In addition, we tested the whole spectrum of sound frequencies between 8 to 32 kHz, and found that mice presented two evident psychological thresholds to deal with this three-choice task (**Fig. 4C5**). Finally, we also implemented sound orientation based d3AFC in another separated group of mice, which actually required longer training duration (45.16±11.46 days) (**Fig. S5B**). The d3AFC was also tested for reversal contingency paradigm and an accelerated learning was revealed (**Fig. S5C**), which potentially provides new insight into the cognitive flexibility ^66^.

Finally, we introduced one of the most challenging cognitive tasks in mouse model, delayed context-dependent attention task, in HABITS **(Fig. 4D)**. This task was previously implemented by light and sound orientation modalities ^67,68^, however, due to the difficulty, it was not well repeated broadly. In HABITS, we tailored this task into a sound-only based but multimodal decision-making task **(Fig. 4D1)**. We constructed this task using three auditory modalities: regular clicks (16 vs. 64 clicks/sec) as context, sound frequency (3k *vs.* 12 kHz) and sound orientation (left vs. right) as the two stimulus modalities. Mice needed to pay attention to one of the modalities, which presented simultaneous during sample epoch, according to the context cue indicated by the clicks (low click rate to sound frequency and high rate to sound orientation), and make a 2AFC decision accordingly. We successfully trained all the 6 mice enrolled in this task, with an average of 48.09 ± 7.54 days **(Fig. 4D2)**. To validate the paradigm’s stability and effectiveness, the direction of stimulus features was presented randomly and independently during the final testing stage.

Mice exceeded criterion performance across different trial types (i.e., unimodal, multimodal w/ conflict, multimodal w/o conflict), indicating effective attention to both stimulus features **(Fig. 4D3)**. Trials with conflicting stimulus features, requiring mice to integrate context information for correct choice, exhibited reduced decision speeds and accuracy compared to trials without conflicting for all tested mice **(Fig. 4D3)**. We further systematically varied the click rate from 16 to 64 Hz to change context contrast. For trials with conflicts, mice decreased their accuracy following the decline of context contrast, formulating a flat psychometric curve, however, for trials without conflicts, mice performed as a near-optimal learner **(Fig. 4D4)**. Meanwhile, as the contrast decreased, mice tended to bias to orientation feature against frequency in conflicting trials, but not for the trials without conflicts **(Fig. 4D5)**. All these results represented mice dealt with different conditioned trials by a dynamic decision strategy synthesizing context-dependent, multimodal integration and perceptual bias. Lastly, the performance of both trial types declined with increased delay duration but maintained criterion above up to 2-sec delay **(Fig. 4D6)**, confirming mice could execute this paradigm robustly in our system.

As a summary, by introducing changes in trial structure, cognition demands, and perceptual modalities, we extended mice behavior patterns in HABITS. These behaviors were usually challenging and very difficult to test previously with manual training. Thus, the training workflow of our system potentially allows for large-scale and efficient validation and iteration of innovative paradigms which aimed to explore unanswered cognitive questions with mouse model.

### Machine learning aided behavioral optimization in HABITS

Mice can be trained to learn challenging tasks in a fully autonomous way in HABITS, however, whether the training efficiency is optimal was unknow. We hypothesized that an optimal train sequence generated by integrating all histories could enhance training procedure, comparing with commonly used random or anti-bias strategies. Benefitted from recent advances in machine teaching (MT) ^38^, and inspired by previous simulations in optimal animal behavior training experiments ^39^, we developed a MT-based automated training sequence generation framework in HABITS to further improve the training qualities.

**Fig. 5A** illustrated the architecture of the MT-based training framework. Initially, mice made an action corresponding the stimuli presented in current trial *t* in HABITS; subsequently, an online logistic regression model was constructed to fit the mice’s history choices by weighted sum of multiple features including current stimulus, history, and rules. This model was deemed as the surrogate of the mouse and was used in the following steps; finally, sampling was performed across the entire trial type repertoire and the fitted model predicted positions of potential future trials in the latent weight space; the trial type with closest position to the goal was selected as the next trials. This entire process forms a closed-loop behavioral training framework, ensuring that the mice’s training direction continually progresses towards the goal.

**Fig. 5.**
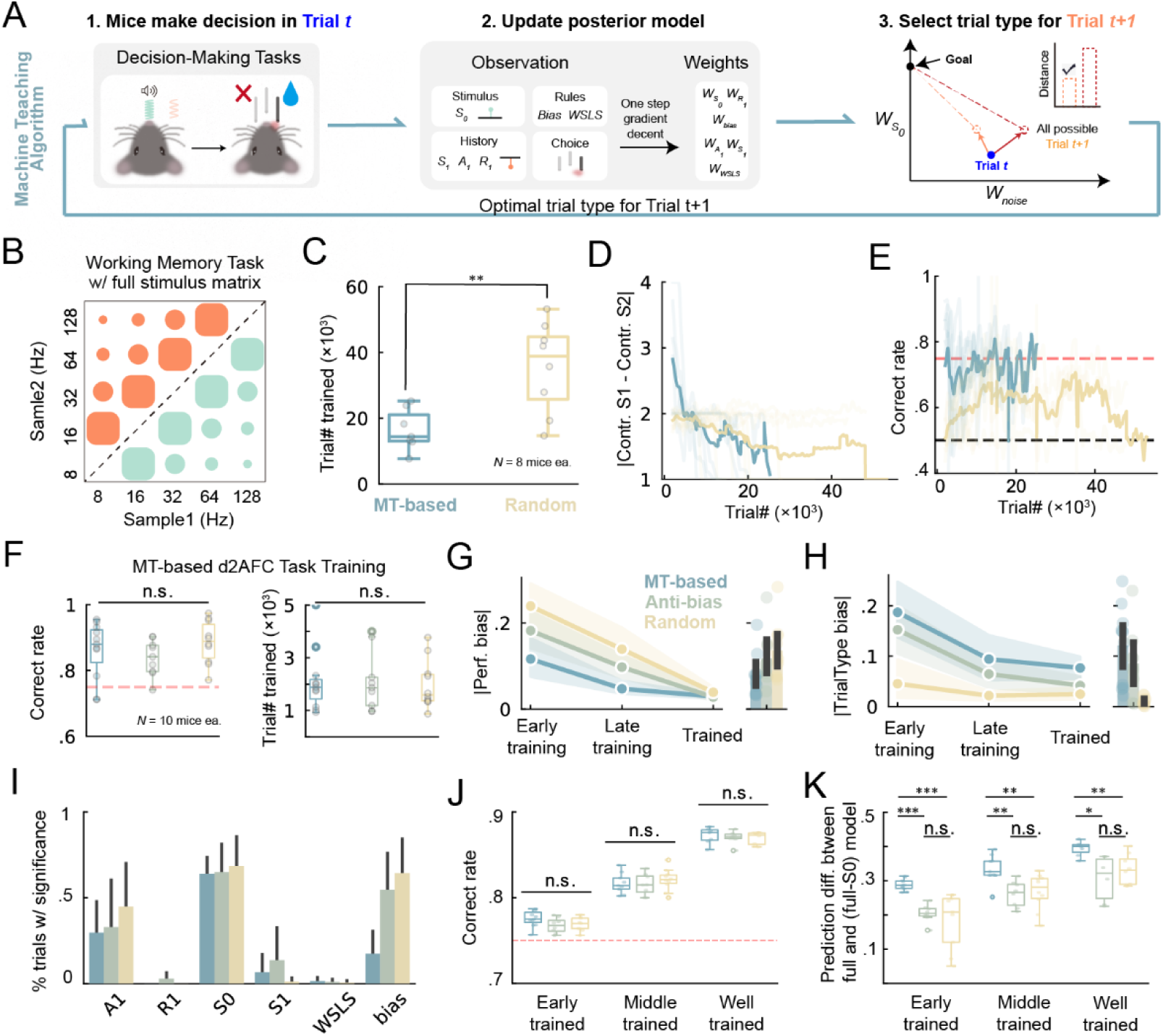
MT enabled faster learning with higher quality. (**A**), The framework of machine teaching (MT) algorithm (see text for details). (**B**), Working memory task as in Fig. 4A, but with full stimulus generation matrix. (**C**), Averaged number of trials needed to reach the criterion for MT-based and random trial type selection strategies. **, p<0.01, two-sided Wilcoxon rank-sum test. (**D**), The absolute difference between contrast (contr.) of sample1 (S1) and sample2 (S2) during training process for the two strategies. (**E**), same as (D) but for correct rate. (**F**), MT-based d2AFC task training. Box plot of correct rate of expert mice (left) and number of trials needed to reach the criterion (right) for different training strategies (MT, anti-bias, and random). n.s., p>0.05, Kruskal–Wallis tests. (**G**), Left, averaged absolute performance bias for the three strategies during different training stages. Right, averaged across training stages. (**H**), same as (G) but for absolute trial type bias. (**I**), Percentage of trials showing significance for different regressors during task learning. (**J-K**), box plot of correct rate (J) and prediction performance difference between the full model and partial model excluding current stimulus (S0) (K) for different trained stage, including early (perf. > 75%), middle (perf. > 80%), and well (perf. > 85%) trained. *, p<0.05, **, p<0.01, ***, p<0.001, n.s., p>0.05, two-sided Wilcoxon rank-sum tests with Bonferroni correction.

We firstly validated the theoretical feasibility and efficiency of the algorithms in simulated 2AFC experiments (**Fig. S6**). Faster increasement in sensitivity to current stimuli was observed through effective suppression of noise like biases and history dependence with MT algorithm **(Fig. S6A-B)**. Notably, if the learner was ideal (i.e., without any noisy), there was no different between random and MT strategies to train (**Fig. S6C**). This implied that the training efficiency was improved by suppression of noise in MT.

To demonstrate MT in real animal training, we initially tested a working memory task similar with Fig. 3B, but with a fully stimulus generation matrix. As shown in **Fig. 5B**, this task utilized a complete set of twenty trial types (colored dots), categorized into four levels of difficulty (dot sizes) according to the distance from decision boundary. Trial type selection using a MT algorithm against a baseline of random selection was tested in two separated groups. Mice trained with the MT algorithm achieved criteria performance with significantly fewer trials comparing with the random group **(Fig. 5C)**; three out of the eight mice in random group even did not reach the criteria performance at the end of training (60 days). We then asked what kind of strategy the algorithm used that supported an accelerated learning. Analysis of the trial type across learning revealed that the MT-based training presented easier trials first then gradually increased the difficulty, i.e., exhibiting a clear curriculum learning trajectory **(Fig. 5D)**. However, this did not mean that the MT only presented easy trials at the beginning; hard trials were occasionally selected when the model deemed that a hard trial could facilitate the learning. This strategy enabled mice to maintain consistently higher performance than random group throughout the training process **(Fig. 5E)**. These results suggested that MT-based method enabled more efficient training for specific challenging tasks.

To further validate the effectiveness of MT in more generalized perceptual decision-making tasks, we trained three groups of mice using random, antibias, and MT strategy respectively in sound-frequency based 2AFC task. Due to the fact that this task was relatively simple, all three groups of mice achieved successful training, with comparable efficiency and final performance **(Fig. 5F)**. But interestingly, MT algorithm effectively reduced mice’s preference towards a specific lickport (i.e., bias) (**Fig. 5G**) throughout the training process by generating trial types with opposite bias more aggressively **(Fig. 5H)**. Using a model-based methodology, we demonstrated that while the MT algorithm minimized bias dependency, it did not increase, and even decreased, mice’s reliance on other noise variables, like previous action *A_1_*, reward *R_1_*and stimulus *S_1_* **(Fig. 5I)**. Notably, we noticed that all trained mice demonstrated similar low bias **(Fig. 5G)**, while only MT algorithm still exhibited relatively high anti-bias strategy during the trained stage **(Fig. 5H)**. This suggests that the MT algorithm might keep regulating cognitive processes actively even in expert mice. To verify this, we segmented the trained trials into early, middle, and well-trained stages based on performance level, and showed that all three groups of mice had similar overall accuracies across stages **(Fig. 5J)**. However, when we examined the reliance on the current stimulus *S_0_*, i.e., to what extend the decision was made according to current stimulus, we found that MT group had significant higher weights for *S_0_* than both anti-bias and random groups **(Fig. 5K)**. This means that MT-generated sequences across all stages encouraged mice to rely only on current stimuli, rather than noise factors. These results suggested that MT-based training had higher quality for both training and trained stage.

In summary, MT algorithm automatically chose optimal trial type sequences to enable faster and more efficient training. By modelling the effect of history, choice and other noise behavioral variables, MT method manifested higher quality training results. Based on these characteristics, the MT algorithm could enhance training efficiency in challenging paradigms and promoted testing robustness in more general paradigms.

### Behavioral optimization for multi-dimensional tasks

One of the advantages of MT algorithm remains that it considers multi-dimensional features altogether and gives the optimal trial sequences to guide the subject to learn. To test this feature, we next expanded the 2AFC task to two stimulus dimensions and trained mice to learn a dynamic stimuli-action contingency. The task was similar to the one applied in Fig. 4D which presented both sound frequency and orientation features but without context cues. As illustrated in **Fig. 6A**, the mice were initially required to focus on sound orientation to obtain reward, while ignoring the frequency (Goal 1). Subsequently, the stimulus-action contingency changed, making sound frequency, rather than orientation, the relevant stimuli dimension for receiving reward (Goal 2). Finally, the relevant cue was still sound frequency but with reversed stimulus-action contingency (Goal 3). Throughout the training process, we employed the MT algorithm to adaptively generate trial types about not only the reward location (left or right) but also the components of the sound stimuli (frequency and orientation combinations), and compared with random control group. MT algorithm allowed for the straightforward construction of a dynamic multi-goal training just by setting the coordinates of target goals within the latent weight space (**Fig. 6B**).

**Fig. 6.**
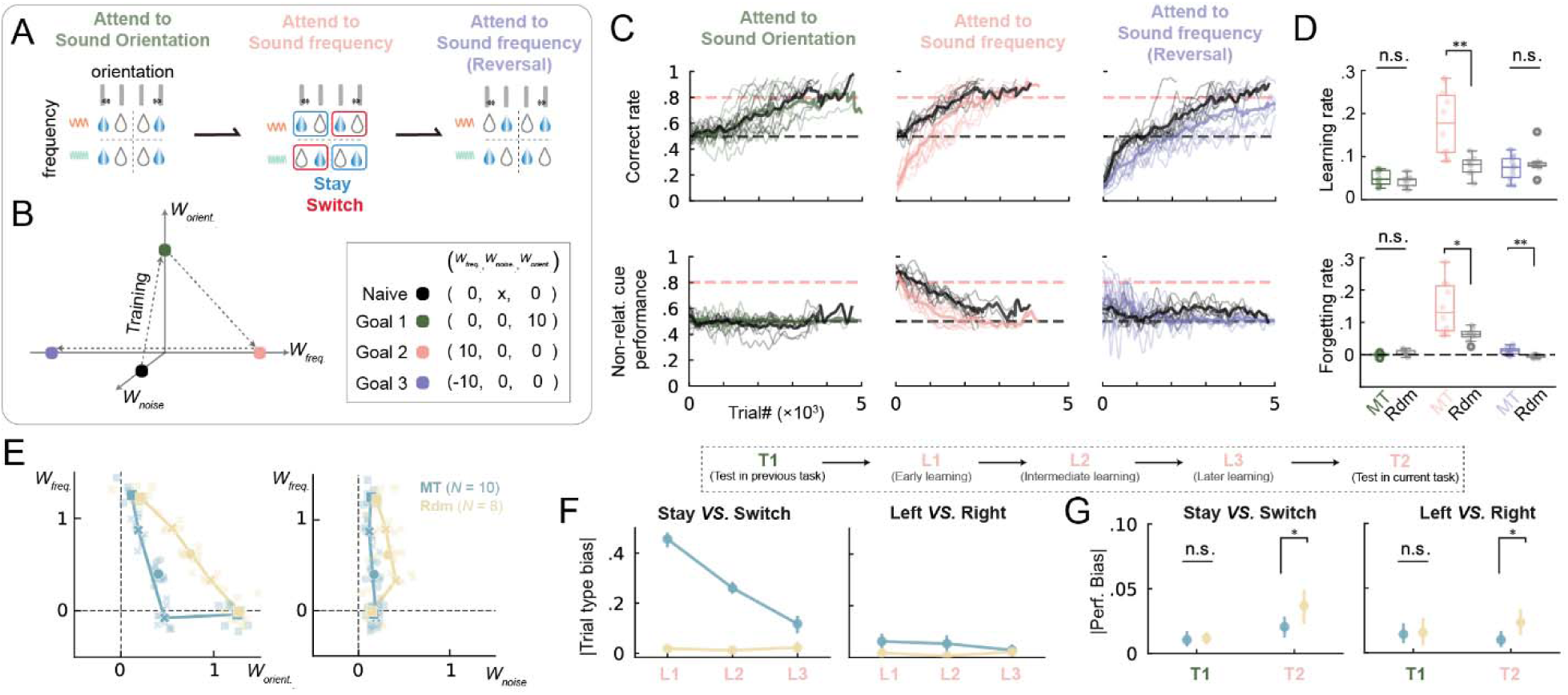
MT manifested distinct learning path with faster forgetting and higher learning rate. (**A**), task structure. (**B**), chart of training path in latent decision space following three goals one by one. (**C**), top, averaged correct rate across grouped mice during training (color, machine teaching; black, random). Bottom, same as top but performance for non-relative cue. (**D**), top, the slopes of linear regression between trial number and correct rate. Bottom, same as top but between trial number and performance for non-relative cue. **, p<0.01; n.s., p>0.05; two-sided Wilcoxon rank-sum tests. (**E**), the learning path of mice (lines) in latent decision space for machine teaching and random training strategies. Light dots represent model weights fitted by individual mice’s behavioral data. Shaded dots, averaged across mice. (Square dots, testing protocol; Cross dots, the first or the last half of trials in learning protocol; Cycle dots, all trials in learning protocol) (**F**), left, averaged absolute trial type bias between stay and switch conditions across grouped mice for the MT and random strategies from L1 to L3. Right, same as middle but for the bias between left and right trials. (**G**), same as (H) but for absolute performance bias in T1 and T2 protocols. L1, the first 500 trials of frequency learning protocol; L2, intermediate trials of frequency learning protocol; L3, the last 200 trials of frequency learning protocol; T1, testing orientation protocol; T2, testing frequency protocol. *, p<0.05; n.s., p>0.05; two-sided t-tests.

Both groups of mice successfully completed all the goals, achieving an overall correct rate of over 80% **(Fig. 6C)**. During the first and the third goal of training, the learning rates of mice in both groups were similar, showing low sensitivity to irrelevant cue. However, in the second goal of training (i.e., transition from orientation to frequency modality), the MT group exhibited a significantly higher learning rate compared to the random group, along with a significantly faster rate of forgetting of the irrelevant cue **(Fig. 6D and S7A-D)**. To construct a paired comparison, we also retrained the MT group mice in the same protocols but with random sequences after forgetting, and confirmed the improvements for both the learning rate and training efficiency (**Fig. S7F-G**). We again employed the logistic regression model to extract the weights for each variable, and plotted the learning trajectories in the latent weight space. MT algorithm manifested distinct learning path against random group in the space (**Fig. 6E**); MT algorithm quickly suppressed the sensitivity of irrelevant modality (i.e., *W_orient_*_._), keeping low sensitivity to noise dimensions (i.e., *W_noise_*) in the meantime. This resulted in a circuitous learning path in the space compared to random group.

We then asked how MT achieved this learning strategy and what is the benefit. Compared to the random sequence, MT algorithm effectively suppressed the mice’s reliance on irrelevant strategies by dynamically adjusting the ratio of stay/switch trials and left/right trial types **(Fig. 6F and S7E)**. After trained, we employed the same random trial sequence to test the performance of both groups. Notably, those mice trained with the MT algorithm exhibited significantly lower left/right bias and stay/switch bias compared to the randomly trained mice **(Fig. 6G and S7H)**. This suggested that the MT algorithm enabled mice to exhibit more stable and task-aligned behavioral training, which implied an internal influence to psychological decision strategies after the MT conditioning.

## Discussion

In this study, we developed a fully autonomous behavioral training system known as HABITS, which facilitates free-moving mice to engage in 24/7 self-triggered training in their home-cage without the need for water restriction. The HABITS equipped with a versatile hardware and software framework, empowering us to swiftly deploy a spectrum of cognitive functions, including motor planning, working memory, confidence, attention, evidence accumulation, multimodal integration, etc. Leveraging the advantages of long-term and parallel running of HABITS, we explored several challenging and novel tasks, some of which introduced new modalities or had never been previously attempted in mouse model. Notably, the acquisition of several tasks spanned more than three months to learn. Benefited from the power of machine teaching algorithms, we endowed the automated training in HABITS with an optimal training sequence which further significantly improved the training efficiency and testing robustness. Furthermore, we extended the machine teaching algorithm to a more generalized form, incorporating multi-dimensional stimuli, which has resulted in diverse training trajectories in the latent space and thus elevated training qualities. Altogether, this study presents a fully autonomous platform that offers innovative and optimal mouse behaviors that could advance the field of behavioral, cognitive and computational neuroscience.

One of the pivotal contributions of this research is provision of an extensive behavioral dataset, derived from ∼300 mice training and testing in more than 20 diverse paradigms. This comprehensive dataset consisted of the entire learning trajectory with well-documented training strategies and behavioral outcomes. This unprecedented scale of data generated in a single study is mainly attributed to the three distinct features of HABITS system. Firstly, the hardware was engineered to fit a broad spectrum of cognitive tasks for mice, diverging from the typical focus on specific tasks in previous studies ^21–34^. It integrated both visual and auditory stimuli across three spatial locations, as well as up to three lickports for behavioral report, offering various combinations to explore. Secondly, the software within HABITS implemented a general-purpose state machine runner for step-by-step universal task learning, and run standalone without PC in the loop, which contrasted with previous system running on PCs ^21,24–26,31^. Thirdly, the cost of the HABITS was quite low (less than $100) comparing previous systems (ranging from $250 to $1500) ^25,26,32,34^, which facilitated large scale deployment and thus high-throughput training and testing. Together, these unique attributes have simplified behavioral exploration, which would otherwise be a time– and labor-intensive endeavor.

Another significant contribution was the expansion of the behavioral repertoire for mice, made possible again by the autonomy of HABITS. We have introduced auditory stimuli and multiple delay epochs into the DMS paradigm ^60–63^, allowing for investigation of working memory across different modality and stages within single trials. Additionally, we have advanced the traditional d2AFC task ^7,43^ to d3AFC, enabling the study of motor planning in multi-classification scenarios. Furthermore, we have implemented delayed context-dependent tasks ^67,68^ based on multi-dimensional auditory stimuli, facilitating research of complex and flexible decision-making processes. Collectively, the advancement of our high-throughput platform was anticipated to improve the experimental efficiency and reproducibility, in either the creation of standardized behavioral datasets for individual paradigms or in the exploration of a multitude of complex behavioral training paradigms.

Last but not the least, we have incorporated machine teaching algorithms into the mouse behavioral training process and significantly enhanced the training efficacy and quality. To our knowledge, this is the first study demonstrating the utility of machine teaching in augmenting animal behavioral training, which complements previous simulation studies ^39^. The impact of machine teaching algorithms is threefold. First, the training duration of complex tasks was substantially reduced, primarily due to the real-time optimization of trial sequences based on mouse performance, which significantly reduces the mice’s reliance on suboptimal strategies. Second, the final training outcomes was demonstrated to be less influenced by the task-irrelevant variables. Prior study has indicated that suboptimal strategies, such as biases, are common among expert mice trained in various paradigms, potentially stemming from their exploration in real-world uncertain environments ^69–72^. Machine teaching-based techniques can significantly reduce the noise dependency, thus facilitating the analysis of the relationship between behavior and neural signals. Third, the machine learning algorithm lowers the barriers for designing effective anti-bias strategies, which were challenging and prone to lopsided in multidimensional tasks. By simply setting the task goals in the fitted decision model, machine teaching can automatically guide the mouse to approach the goal optimally and robustly.

Our study was designed to standardize behavior for the precise interrogation of neural mechanisms, specifically addressing within-subject questions. However, investigators are often interested in between-subject differences—such as sex differences or genetic variants—which can have long-term behavioral and cognitive implications ^73,74^. This is particularly relevant in mouse models due to their genetic tractability ^75^. Although our primary focus was not on between-subject differences, the dataset we generated provides preliminary evidence for such investigations. Several behavioral readouts revealed individual variability among mice, including large disparities in learning rates across individuals (Fig. 2I), differences in overall learning rates between male and female subjects (Fig. 2D vs. Fig. S2G), variations in nocturnal behavioral patterns (Fig. 2K), etc. Furthermore, a detailed logistic regression analysis dissected the strategies mice employed during training (Fig. S2B). Notably, the regression identified variables associated with individual task-performance strategies (Fig. S2C), which also differed between manually and autonomously trained groups (Fig. S2D). Thus, our system could facilitate high-throughput behavioral studies exploring between-subject differences in the future.

Our study marks the inaugural endeavor to innovate mouse behavior through autonomous setups, yet it comes with several limitations. Firstly, our experiments were confined to single-housed mice, which is known to influence murine behavior and physiology, potentially affecting social interaction and stress levels ^76^. In our study, individual housing was necessary to ensure precise behavioral tracking, eliminate competitive interactions during task performance, and maintain consistent training schedules without disruptions from cage-mate disturbances. However, the potential of group-housed training has been explored with technologies such as RFID ^28,29,32–34^ to distinguish individual mice, which potentially improving the training efficiency and facilitating research of social behaviors ^77^. Notably, it has shown that simultaneous training of group-housed mice, without individual differentiation, can still achieve criterion performance ^25^. Secondly, we have not yet analyzed any videos or neural signals from mice trained in the home-cage environment. Recent studies have harnessed a variety of technologies and methodologies to gain a deeper understanding of natural animal behavior in home-cage environments^78,79^. Voluntary head-fixation, employed in previous studies, has facilitated real-time brain imaging ^17,18,21,29,30^. Future integration of commonly-used tethered, wireless head-mounted ^80^, or fully implantable devices ^81,82^, could allow for investigation of neural activity during the whole period in home-cage. Lastly, while HABITS achieves criterion performance in a similar or even shorter overall days compared to manual training, it requires more trials to reach the same learning criterion (Fig. 2G). We hypothesize that this difference in trial efficiency may stem from the constrained engagement duration imposed by the experimenter in manual training, which could compel mice to focus more intensely on task execution, resulting in less trial omissions (Fig. 2F). In contrast, the self-paced nature of autonomous training may permit greater variability in attentional engagement ^83^ and inter-trial-intervals, which could be problematic for data analysis relaying on consistent intervals and/or engagements. Future studies should explore how controlled contextual constraints enhance learning efficiency and whether incorporating such measures into HABITS could optimize its performance.

The large-scale autonomous training system we proposed can be readily integrated into current fundamental neuroscience research, offering novel behavioral paradigms, extensive datasets on mouse behavior and learning, and a large cohort of mice trained on specific tasks for further neural analysis. Additionally, our research provides a potential platform for testing computational models of cognitive learning, contributing to the field of computational neuroscience.

## Materials and Methods

### Design and implementation of HABITS

#### Architecture

A single HABITS was comprised of a custom home-cage and integrated behavioral apparatuses. All the building materials were listed in the supplementary Table I, with source and price information provided. The home-cage was made of acrylic panels, with a dimension of 20 × 20 × 30 cm^3^. Top panel was movable and could equipped with cameras to record mouse natural behaviors. A compatible tray was located at the bottom of the home-cage, facilitating bedding materials changing. A notch was designed in the front of tray where an elevated platform was installed. The platform formed an arch-shape to loosely constrain the mouse body when the mouse stepped on it to perform task. A micro load cell was installed beneath the platform and used for daily body weighting.

Most of the behavioral apparatuses were installed in the front panel of the home-cage. A lickport holder with up to seven slots was installed in front of the weighting platform. Three lickports (1.5 mm diameter, 10 mm apart) were used in this study. Water was drawn by peristaltic pumps from water tanks (centrifuge tube, 50 ml) to the lickports. Three groups of LEDs and buzzers for light and sound stimuli were extruded from the front panel and placed in the left, right and top position around the weighting platform. Notably, the top module contained a RGB LED, but white LEDs for the others. Buzzers were the same in all stimulus modules and produced 3-15 kHz pure tones at 80 dB. In some experiments (Fig. 3E, Fig. 4C), the top buzzer was replaced with a micro ultrasound speaker (Elecfans Inc.) which was able to emit 40khz pure tone for up to100 dB.

#### Control system

The core of the control system was a microprocessor (Teensy 3.6) which interacted with all peripheral devices and coordinated the training processes (Fig. S1A). The microprocessor generated PWM signal to directly control the sound and light stimuli. Reward water was dispersed by sending pulses to solid state relays which controlled the pumps. Two toggle switches were used for flushing the tubing. Each lickports were electrically connected to a capacitive sensing circuit for lick detection. Additionally, another switch was used for manually controlling the start and pause of training process. Real-time weight data were read from the load cell at a sampling rate of 1 Hz. A Wi-Fi module was connected with the microprocessor to transmitted data wirelessly to a host computer. Meanwhile, all the data were also stored on a local SD card, with the microprocessor’s clock as the timestamps for all behavioral events.

We have developed a software framework for constructing behavioral training programs, which is a general-purpose state machine runner for training animal behaviors (gpSMART, https://github.com/Yaoyao-Hao/gpSMART). This framework supported construction of arbitrarily complex cognitive behavioral paradigms as state machines (Fig. S1B). Basically, each state was comprised of a unique name, output actions, transition conditions and maximum timing. Within each trial, the microprocessor generates the state matrix based on the defined state machine and executes the state transition according to the external events (e.g., licks) or timing (e.g., a delay period of 1.2 sec). This is similar to the commonly used Bpod system (Sanworks Inc.), but gpSMART could run on microprocessors with a hardware-level time resolution. Between trials, training protocol updating, behavioral data recording, and wireless data communication were executed. Various training assistances (e.g., free reward) were also performed when necessary to help the training processes. All the training progress and protocols were stored on SD card for each mouse; thus, training can be resumed after any pause event and supports seamless switching between multiple HABITS systems. The system was designed to operate standalone without PC connected. Finally, the firmware on the microprocessor could be updated wirelessly to facilitate paradigm changing.

#### High-throughput training and GUI

We constructed over a hundred of HABITS to facilitate large-scale, fully autonomous in-cage behavioral training (Fig. S1D). Each HABITS was piled on standard mouse cage racks, with sound-proof foams installed between them to minimize cross-cage auditory interference. The cage operated independently with each other, with only a 12V standard power supply connected. The training room were maintained under a standard 12:12 light-dark cycle. All the cages communicated with a single PC via unique IP addresses using the UDP protocol.

To monitor the training process of all the cages, a MATLAB-based graphic user interface (GUI) running on a PC was developed (Fig. S1C). The GUI displayed essential information for each mouse, such as the paradigm name, training duration, training progress, and performance metrics like long-term task accuracy, weight changes, and daily trial numbers, etc. Meanwhile, the whole history of training performance, detailed trial outcomes in the last 24-hour and real-time body weight could be plotted. The GUI also enabled real-time updating of each cage’s training parameters for occasional manual adjusting. Training settings can also be modified by physically or remotely updating the SD card files.

### Mice and overall training procedures

#### Mice

All experimental data used in this study were collected from a total of 302 mice (C57BL/6J). For most of the autonomous experiments, males were used with starting age at around 8-week (see Table I). A separate group of 6 females were tested in a sound-frequency based 2AFC task (**Fig. S2G**). Mice were single housed in our home-cage systems for ranging from 1 to more than 3 months. A group of 6 mice were used for supervised manual training. Another 6 mice were used for *ad libitum* reward testing in HABITS. All experiments were conducted in compliance with the Zhejiang University Animal Care Committee’s regulations.

#### Workflow for behavioral testing in HABITS

The entire workflow for fully automated behavioral training experiment in HABITS can be divided into three stages (**Fig. S1E**). The first stage was the initialization of HABITS. This involved setting up the home cage by placing an appropriate amount of bedding, food, cotton, and enrichments into the drawer of the home cage. Behavioral paradigms and training protocols, programmed within our software framework, were then deployed on the microcontroller of HABITS. Each mouse was provided with a unique SD card that stores its specific behavioral training data, including the training paradigm, cage number, initial paradigm parameters, and progresses. The load cell was initialized through the host computer, which includes zeroing and calibration processes. The flush switch of the peristaltic pump was activated to fill the tubing with water. Finally, the mouse was placed into the HABITS after initial body weight measuring. Note that any of habituations or water restriction were not required.

The second stage was the fully autonomous training phase, during which no intervention from the experimenter was needed. Typically, this stage included three main training sub-protocols: habituation, training, and testing. During the habituation phase, free rewards are randomly given on either lickports to guide the mouse establish connection between lickports and water reward. Subsequently, in the training phase, the protocols are gradually advanced, form very easy one to the final paradigm, based on the learning performance of the mouse. Assistances, like reward at correct lickport, were gradually decreased as the mouse learned the task. Finally, predefined behavioral tests, such as psychometric curve testing, random trials, additional delays, etc. were conducted. The entire training process of all cages were remotely monitored via the GUI. The bedding in the drawer were replaced every other week to ensure that the mouse lives in a clean environment.

The third stage involved data collection and analysis. All raw data, including detailed event, trail and configuration information, were stored on the SD card; data wirelessly transmitted to PC were mainly used for monitoring. These behavioral data were analyzed offline with Python and the mice were ready for other subsequent testing.

#### Manual Training

To compare with fully autonomous training, we also used HABITS as a behavioral chamber to perform manual training protocol for freely moving mice. The mice were first single-housed in standard home cages and subjected to water restriction. After several days, when the mice’s body weight dropped to approximately 85% of their original weight ^9^, behavioral training began. The mice were trained in a session-based manner; in each session, experimenters transferred the mice from the standard home-cage to HABITS, where they underwent 1-3 hours of training to receive around 1 ml of water. The amount of water consumed was estimated by HABITS based on the number of rewards. HABITS weighed the mice daily, ensuring that all mice maintained stable body weight throughout the training process. The manually trained mice underwent the same training protocols as in the autonomous ones (**Fig. S2A**). Once the mice completed the final protocol and reached the criterion performance (75%), they were considered successfully trained. After completing the manual training, the mice were then transitioned into autonomous testing in HABITS (**Fig. S2E**).

### Behavioral data analysis

#### Bias calculation

We calculated mice’s bias toward different trial types, e.g., left and right, by evaluating their performance under these trial types (perf. bias). The strength of the bias was quantified by calculating the absolute difference between the proportion of performance under specific trial type relative to the summed value across trial types, and the balance point, i.e., 50%. Similarly, we applied this method for presentation of trial sequence to compute the trial type bias during paradigm training, illustrating the dynamic changes in training strategies.

#### Data preprocessing

For the fully autonomous training, we excluded data from the habituation phase, as we believed the mice had not yet understood the structure of the trials during that stage. Additionally, we removed trials where the mice did not make a choice, i.e., no-response trials. For each mouse, the trials were concatenated in chronological order, ignoring the time span between trials during the continuous multi-day home-cage training sessions; the same approach was applied to manual training data. We then organized the data for each mouse into multiple 500-trial windows, sliding from the beginning to the end of the training with a step size of 100-trial. Windows containing fewer than 500 trials at the end of the dataset were discarded. We assumed that within each window, the mouse employed a consistent strategy, and a new logistic regression model was fit in each window.

#### Logistic regression of behavioral data

Similar to our previous study ^27^, we employed an offline logistic regression model to predict the choices made by the mice (**Fig. S2B-D, Fig. 5I**, **Fig. 6**). This model calculates a weighted sum of all behavioral variables and transforms the decision variable into a probability value between 0 and 1 using a sigmoid function, representing the probability of choosing the left side. The variables include the current stimulus (*S_0_*; –1 for licking left trials; 1 for licking right trials), the previous stimulus (*S_1_*), reward (*R_1_*; –1 for no reward; 1 for reward), action (*A_1_*; –1 for left choice; 1 for right choice), win-stay-loss-switch (*WSLS*; which is *A_1_*× *R_1_*), and a constant bias term (*bias*). The model can be formulated by the following equation:

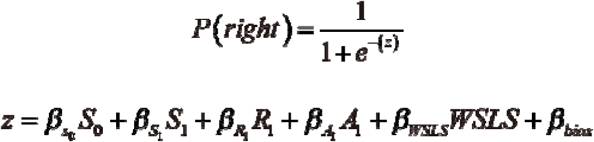

where the β’s were the weights for the regressors. We used 0.5 as the decision threshold: predictions above 0.5 were classified as right choices, while below were classified as left choices.

Model performance was assessed using 10-fold cross-validation. For each cross-validation iteration, 450 trials were randomly selected as the training set, and gradient descent was employed to minimize the cross-entropy loss function. The remaining 50 trials were used as the test set. Training was considered complete (early stopping) once the calculated loss in the test set stabilized. The accuracy of the model in predicting the mouse’s choices in the test set was recorded as the result of one cross-validation iteration. This process was repeated 10 times, and the final performance of the model was averaged across all iterations.

#### Significance calculation

To evaluate the contribution of each regressor, we compared the performance of a partial model, where a specific variable was removed, with that of the full model. Specifically, the value of the variable in testing was set to zero, and we checked whether the performance of the partial model showed a significant decline. We applied a corrected t-test using a 10×10 cross-validation model comparison method to compute the *p*-value ^84^. For each window, we trained 100 models, and the performance differences between the partial and full models formed a *t*-distribution. By examining the distribution of performance differences, we determined the significance level of each regression variable’s contribution. When *p* < 0.05, the regression variable was considered to have a significant contribution to predicting the mouse’s choice in that window. Additionally, we calculated the proportion of windows across the entire training process in which a particular regression variable had a significant contribution, to estimate the degree to which the mouse relied on that variable. The same significance evaluation method was applied to both autonomous and manual training, allowing for direct comparison of the learning strategies employed in two conditions at the individual mouse level (**Fig. S2B-D, Fig. 5I**).

#### Logistic regression in evidence accumulation: Multimodal integration

For each mouse in evidence accumulation task (**Fig. 3 D4**), we trained a group of logistic regression models to estimate the psychophysical kernel. The entire sample period was divided into 25 bins of 40 ms each, with each bin assigned a weight to predict the mouse’s choice. An event occurring in a bin was set to 1, otherwise, set to 0. We trained 100 pairs of models for each mouse. Each pair of models trained using 10% total trials randomly. Each pair of models included one model trained on the original data and another trained on data with bin-wised shuffled within each trial. The psychophysical kernel for each mouse was derived by averaging the first 100 models, compared to a baseline kernel obtained from the second. Finally, we averaged the results across all 13 mice to statistically estimate the temporal dynamics of mice’s evidence dependence in this task.

### Behavioral tasks and training protocols

#### General training methodology

In the fully autonomous behavioral training process, all mice learn the required behavioral patterns through trials and errors. The training protocols were pre-defined based on experience. Given that the entire training process is long-term and continuous, a free reward is triggered if a mouse fails to obtain water within last 3-or 6-hour period, ensuring the mouse receives sufficient hydration. Throughout the training process, we employed a custom-designed ‘anti-bias’ algorithm to avoid mice always lick one side. Basically, we implemented several priority-based constraints to prevent mice from developing a preference for a particular reward direction:

– *Highest Priority*: If a mouse consecutively made three errors or no-response in a specific trial type, the next trial would maintain the same reward direction.
– *Second Priority*: If three consecutive trials shared the same reward direction (including no-response trials), the reward direction would switch in the subsequent trial.
– *Third Priority*: The reward direction of the next trial was sampled based on the average performance of left and right trials, using the following formula:

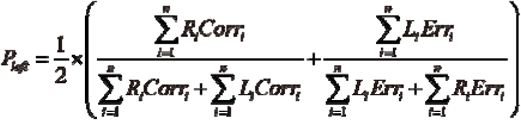

where *N* represented the number of recorded historical trials (set to 50 in our case). *Ri* and *Li* were set to 1 if the reward direction of the *i_th_* historical trial was right or left, respectively; otherwise, they were set to 0. Similarly, *Corr_i_*and *Err_i_* were set to 1 or –1 if the mouse’s choice in the *i_th_* trial was correct or incorrect, respectively; otherwise, they were set to 0. *P_left_* represented the probability of left trial type in the next trial.

#### d2AFC with multi-modal

The task of d2AFC, delayed two-alternative forced choice, required mice to learn the stimulus-action contingency separated by a delay for motor planning. A complete trial consisted of three parts: the sample, delay, and response epochs. The sample epoch lasted for 1.2 sec and is accompanied by auditory or visual stimuli. The delay epoch elapsed for another 1.2 sec, during which mice were required to withhold licking until a 100 ms response cue (6 kHz tone) was played. Any premature licking (early licks) during this period immediately paused the trial for 300ms. Response epoch lasted for 1 sec. The first lick made by the mouse during this period was recorded as its choice for the current trial, and feedback is provided accordingly. A correct lick delivered approximately 0.25µl of water to the corresponding spout (achieved by activating the peristaltic pump for 30ms), while an incorrect choice results in an immediate 500 ms white noise and 8000 ms timeout for penalty. After each trial, the mouse must refrain from licking for 1000 ms before the next trial began automatically. If the mouse failed to make a choice during the response period, the trial was marked as a no-response trial, and the mouse must lick either spout to initiate the next trial. The stimuli modalities tested in this study were as follows:

– Sound frequency modality: A 3 kHz tone corresponds to a left choice, while a 10 kHz tone corresponds to a right choice.
– Sound orientation modality: A sound from the left speaker corresponds to a left choice, while a sound from the right speaker corresponds to a right choice.
– Light orientation modality: The left white LED lighting up corresponds to a left choice, and the right white LED lighting up corresponds to a right choice.
– Light color modality: The top tricolor LED lighting up blue corresponds to a left choice, and red corresponds to a right choice. For light color modality, we tested multiple variations since the mouse did not learn the task very well, including green *vs.* blue and flashed green *vs.* blue.

In the reaction time version of the paradigm (RT task, Fig. S3), a central spout was introduced in addition to the left and right lickports, to allow the mouse to self-initiate a trial. The mouse must lick the central spout to initiate a trial; licking either side following would result in a brief (100ms) white noise and immediate termination of the trial, followed by a timeout period. During the sample epoch, a tone from the top speaker (3 kHz for left reward; 10 kHz for right reward) plays for 1000ms. The mouse can immediately indicate its choice by licking either side lickports, which terminated the sample period and triggered trial outcomes as above. An inter-trial interval (ITI) of 1000ms was followed. The next trial required the mouse to lick the central spout again. The central spout did not provide any rewards; all rewards are contingent upon the mouse’s choice of the left or right spouts.

#### Training and testing of other tasks

The training and testing method for all other tasks were detailed in the text of supplementary materials.

### Machine teaching algorithms

We employed machine teaching (MT) algorithm ^41^ to design an optimal trial sequence that enables the mice to rapidly approach a target state, i.e., trained in a specific task. In this context, the MT algorithm can be referred to as the ‘teacher’ while the mice are the ‘students’; the size of the training dataset is termed ‘teaching dimension’ of the student model ^38^. Specifically, the teacher samples from a pre-defined discrete dataset, and the student updates its internal model using the sampled data. The teacher then prepares the next round of training based on the student’s progress, creating a closed-loop system. In this study, a logistic regression model was employed to infer the internal decision-making model of the mice based on their choices trial-by-trial (i.e., model-based). The model was updated in real-time and used for the optimization and sampling of subsequent trials. L1 regularization and momentum were introduced to smooth the fitted weights, mitigating overfitting and oscillations. The mice’s choices and outcomes served as feedback for MT. Figure 5A illustrated the complete closed-loop optimization process of MT algorithm. It was similar to an imitation teacher ^41^ whose objective was to iteratively minimize the distance between the model weights of next trial and target.

In detail, the logistic regression model to fit the choices of mouse was updated trial-by-trial according to the following formula:

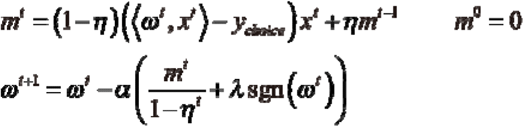

where the parameter ω*^t^* represents the decision model parameters fitted to the current and past choices (i.e., *y_choice_*) performed by the mice at the *t*-th trial. The hyperparameter λ controls the strength of L1 regularization. Momentum parameter η determines the window width for exponential smoothing of the loss gradient. *m* represents an exponential smoothed gradient across past trials.

Then, the objective of this algorithm can be formalized as the following equation:

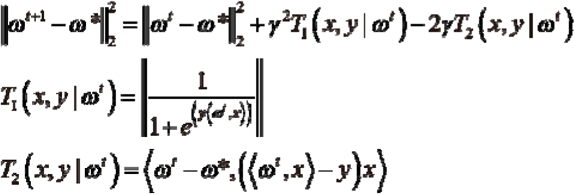

where, (*x, y*) represents a pair of stimuli-action contingency. The parameter ω*\**denotes the target weight within the implicit decision space of the simulated mouse model, typically set to converge to the model weights according to the current task rules. *T_1_* can be interpreted as the trial difficulty, predicting the probability of incorrect choices performed by mice in this trial, and *T_2_* as the effectiveness of this trial, predicting the correlation between the upcoming mouse behavioral strategy updates and the shortest learning path between ω*^t^* and ω*\**. The balance between these two metrics is achieved through hyperparameters γ, i.e., hypothetical learning rate of mouse. In this study, we assume that this model reflects the actual decision-making process of the mice. Subsequently, this algorithm can select the next trial type using the following formula:

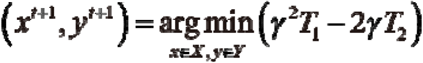

where *X* and *Y* represent the repertoires of available trial types and action, respectively.

Finally, we presented the selected stimuli *x* in the next trials and expected the mouse made correct/incorrect choice *y* and received reward/punishment.

We have implemented the aforementioned MT-based optimization algorithm on the microprocessors of HABITS, as a superior alternative to existing ‘anti-bias’ algorithms. The computation load for running MT optimization on the microprocessors was high and latency was relative long comparing trial resolution. However, since the computation was conducted between trials, it did not interfere with the execution of the trials themselves under the gpSMART framework. Additionally, for each mouse, additional files were used to record the hyperparameters and weight changes of the online logistic regression model for each trial. No-response trials were not used for model fitting. However, if the previous trial was a no-response trial, the history-dependance regressors of current trial were set to 0.

#### Simulation experiments

We employed a logistic regression model as the student, tasked with completing 2AFC task, which involved mapping the current stimulus *S_0_*to a choice while ignoring interference from other features. Another logistic regression model, serving as the imitation teacher, was then used to fit the choices made by the student. Both models operated within the same feature space and utilized the same update algorithm. The hyperparameters were set as follows: α=0.1, η=0.9, γ=1, and λ=0.1.

We first simulated the biases and history-dependence typically observed in naïve mice during the early stages of training by setting the initial values of *S_1_* and *bias* to –2 and 2, respectively. During the trial-by-trial update process, we tracked the changes in the student’s weights under the MT algorithm and compared them to those under random training, which served as the baseline. Additionally, we simulated conditions without noise to further examine the differences between the MT algorithm and random training in influencing the student’s weight updates.

#### Animal experiments 1: Working Memory Task with Full SGM

The first task we tested with MT algorithm was working memory task with full 5×5 stimulus matrix (Fig. 5B). The delay duration was set to 500ms. Mice were randomly assigned to two groups: random (*N* = 8) and MT (*N* = 7). Once the mice achieved a 75% correct response rate in the eight most challenging trial types, they advanced to the testing phase. At this stage, only the eight most challenging trial types were randomly presented. When the correct rate reached 75%, the mice are considered to have learned the task. MT used seven features as the inputs for the logistic regression model: *bias*, *S_0_* (the frequency of the first stimulus, where {8, 16, 32, 64, 128} Hz corresponds to values of {-1, –0.5, 0, 0.5, 1}), *T_0_* (the frequency of the second stimulus), *S_1_*, *A_1_*, *R_1_*, and *WSLS*. Hyperparameters were set as follows: α=0.01, η=0, γ=0.03, and λ=0.

#### Animal experiments 2: 2AFC task

In 2AFC task (Fig. 5F), Mice were divided into three groups, each using different methods to generate the stimulus frequency (3 kHz or 10 kHz) for the next trial: random selection (*N* = 10), the anti-bias strategy (*N* = 10), and the MT algorithm (*N* = 10). The online logistic regression model settings for this task remained consistent with those in Figure S2. The hyperparameters for the MT algorithm were set as follows: α=0.1, η=0.9, γ=1, and λ=0.1.

#### Animal experiments 3: dynamic 2AFC task with multi-dimensional stimuli

In the third task, we expanded the stimulus dimension of 2AFC with the combination of sound frequency (high and low) and orientation (left and right), to test MT algorithm (Fig. 6A). In each trial, the stimulus could be one of the four combinations, but whether the mouse should attend to frequency or orientation modality was not informed. In other words, no any modality cues were presented for mice and mice must rely solely on feedback from past trials to identify what the current modality was. Mice were divided into random (*N* = 8) and MT groups (*N* = 10). In MT group, both the stimulus combinations and reward direction are determined by the MT algorithm simultaneously. The entire task involved learning three different rules one by one, including sound orientation, sound frequency, and reversed sound frequency. When the mice in each group achieved an accuracy of 80% in the first 500 choice trials under the current rule, they advance to a testing protocol consisting of at least 100 random trials. When a mouse achieved 80% accuracy in the last 100 trials of the testing protocol, it transitioned to next rule ^85^. Hyperparameters were set as follows: α=0.1, η=0.9, γ=1, and λ=0.1.

## Statistics

To maximize the utility of HABITS in a wider range of paradigms, we usually employed 6 mice per paradigm. For experiments where we aimed to conduct between-group comparisons, we increased the sample size to 10 to ensure the stability and reliability of statistical significance. All significance tests were conducted by comparing different groups of animals (e.g., comparing performance levels across different mouse groups). Non-parametric tests, such as the Wilcoxon signed-rank test or rank-sum test, were used for comparisons between two groups, and the Kruskal–Wallis test was used for comparisons among three groups, unless otherwise stated in the figure legends. Data are presented as mean ± 0.95 confidence intervals (CI) across animals, as specified in the figure legends. In the box plots, the center lines represent the median values, the box limits indicate the upper and lower quartiles, the whiskers show 1.5 times the interquartile range, and the points represent outliers. Significance levels are denoted as *, p < 0.05, **, p < 0.01, and ***, p < 0.001 in all figures. All data analyses were performed using Python (version 3.8.13) with the following packages: NumPy (1.21.5), SciPy (1.10.1), matplotlib-inline (0.1.3), pandas (1.3.4), PyTorch (1.11.0), and seaborn (0.13.0).

## Supporting information

old Supplemental video 1

old Supplemental video 2

## Author contributions

Conceptualization: BY, YH; Methodology: BY, KX, YW, YH; Investigation: BY, PL, HX, YH; Visualization: BY, YH; Supervision: KX, YW, YH; Writing—original draft: BY, YH; Writing—review & editing: BY, KX, YW, YH.

## Data and materials availability

The guidance for construction of HABITS, all the training programs, and example behavioral data are available on GitHub (https://github.com/Yaoyao-Hao/HABITS). The general-purpose state machine runner for training animal behaviors (gpSMART) is also available on GitHub (https://github.com/Yaoyao-Hao/gpSMART). All data in the main text or the supplementary materials are available on https://doi.org/10.6084/m9.figshare.27192897.

## Supplementary Materials

### Tasks tested in HABITS

#### Value-based dynamic foraging task

In Figure S4, we introduced the dynamic foraging task. Mice must first lick the center spout to initiate a trial block. Each block consists of one or more trials, and if a mouse fails to make a choice in any given trial, the block ends and waits for the mouse to trigger the next one. Within a trial block, trials are separated by an inter-trial interval (ITI) ranging randomly from 0.5 to 2 seconds, with an average of 1 second. The next trial automatically begins after the ITI.

At the start of each trial, a 500ms light cue is emitted from the top LED, signaling the beginning of the trial. This is followed by a delay period lasting either 1 or 1.5 seconds, during which the mouse must restrict from licking the left or right spouts. If the mouse licks prematurely, the trial pauses for 300ms. After delay, a 6 kHz pure tone lasting up to 2000ms is played from the top buzzer, prompting the mouse to choose by licking either the left or right spout. The first lick during this period is registered as choice for this trial, followed by an auditory cue lasting 50ms (3 kHz for left, 10 kHz for right). If the choice matches the assigned reward direction, a 0.25μL water reward is dispensed from the corresponding spout.

In this task, mice are unable to obtain any useful cues from single trial to make a rewarded choice. All rewards in individual trials are randomly assigned to the left or right spouts according to default probabilities. We use a minimum of 500 trials per rotation, during which the reward probabilities for left and right spouts are rotated in the following sequence: {60:10}, {52.5:17.5}, {10:60}, and {17.5:52.5}. Additionally, mice cannot obtain rewards from one spout if the reward allocated to the other spout has not yet been consumed. Only when the mouse chooses the spout with the higher reward probability in more than 50% probabilities, the rotation advances. As the mouse’s performance hits criteria, the frequency of rotations increases, starting from minimum every 500 trials and gradually decreasing to every 100 trials.

#### Contingency reversal learning

In Figure 3A, we extended the self-initiated paradigm (as shown in Fig. S3) by introducing multiple reversals in the relationship between stimulus and action. Specifically, when a mouse achieves an accuracy rate of over 75% in the most recent 100 trials, the association between sound frequency and reward direction is immediately reversed, while the trial structure and parameters remain unchanged. The mouse receives no additional cues indicating the rule reversal, relying solely on reward and error feedback to adapt. Subsequently, the contingency is reversed each time the mouse reaches the specified accuracy under the current rule.

#### Working Memory Task

In Figure 3B, the mouse begins by licking the central spout to initiate a trial. After a brief delay, the first stimulus is presented, lasting 500ms, and consisting of a sequence of regular clicks. This is followed by a 200ms delay period, during which the mouse must refrain from licking the side spouts. Any licking during this delay immediately terminates the trial and results in a noise burst and a timeout penalty. Following the delay, a second 500ms stimulus of regular clicks is presented. The mouse must then wait for a brief response cue after the delay and indicate its choice by licking either side spout, receiving feedback based on its decision: correct choices yield a water reward, while incorrect choices trigger a 500ms noise burst and a timeout. The correct choice is determined by comparing the click rates of the two stimuli—when the first stimulus has a higher click rate, the reward is on the right side, and when it is lower, the reward is on the left.

The trial types are generated using an anti-bias algorithm to determine the left or right reward direction, after which one of the four possible click rates for that direction is randomly selected as the stimulus for the current trial. The SGM comprises five click rates separated by octave intervals: 8, 16, 32, 64, and 128 Hz.

During the testing phase, the delay period is randomly sampled between 500 and 2000ms. Additionally, in half of the trials, the click rate of the second stimulus is fixed at 32 Hz, while the click rate of the first stimulus is randomly sampled from the set {16, 20, 25, 32, 40, 50, 64 Hz}. If the first click rate is set at 32 Hz, the rewarded choice is randomly assigned to the left or right spout.

In Figure S5, a 3×3 SGM was used with a two-octave difference between the click rates of the first and second stimuli. After training, a certain percentage of probe trials were introduced, where the click rates of the two stimuli were identical, and the reward direction was randomly assigned. The mouse’s performance in these probe trials was at chance level, indicating that the mouse indeed relied on comparing the frequencies of the two stimuli to complete the task.

#### Evidence accumulation

In Fig. 3C, a new trial block is initiated when a mouse licks either of the two spouts. After a brief delay (200-500ms), independent click sequences, following an exponential distribution with predefined parameters, are generated from speakers on both sides. After a second brief delay period, a response cue signals the mouse to choose by licking either the left or right spout. The mouse should select the left spout if the click rate on the left is higher, and the right spout if the click rate on the right is higher. Correct choices are immediately rewarded with 0.25 µl of water at the corresponding spout, while incorrect choices result in a noise burst and a timeout penalty.

The 1000ms sample period is divided into 40 bins, each lasting 25ms. To approximate a discrete event sequence with an exponential distribution, a coin-flip method is used for each bin to determine whether a click event occurs, based on the set probability. Each click event consists of a 10ms, 10 kHz auditory cue followed by a 15ms interval. The hazard rates for generating clicks on the left and right sides are randomly selected from {39:1, 37:3, 31:9, 26:14}. The onset time of the clicks is randomly chosen from within the first 0 to 900ms of the sample period, in 100ms increments.

#### Evidence accumulation with multimodal integration

In Fig. 3D, a trial block is initiated when the mouse licks either the left or right spout. After a brief delay, a top speaker emits click sequences following an exponential distribution, while both the left and right white LEDs flash according to the same event sequence. Following the delay period, a response cue indicates the start of the response period, during which the mouse must decide if the event rate exceeds 12 clicks per second. If the event rate is below 12, the mouse should choose the left spout; if it is above 12, the right spout should be chosen. For trials where the event rate is exactly 12, the reward is randomly assigned to either the left or right spout.

The sample period is divided into 40 bins, each lasting 40ms. Similar to the previous task, a coin-flip method is used to approximate the exponential distribution of discrete event sequences, determining whether a multimodal event occurs in each bin. Each multimodal event consists of 20ms light flash paired with an auditory click, followed by 20ms interval. For trials where the reward is assigned to the left spout, the hazard rate for event sequence generation is 4; for trials with the reward on the right spout, the hazard rate is 20. The sample period lasts for 1600ms, with the event onset fixed at 600ms to ensure that the events continue for 1000ms.

During the testing phase, a portion of the trials had event onset times randomly selected from 0, 400, 800, or 1200ms after the start. We also tested the mouse’s performance under unimodal conditions, using only clicks or only flashes as stimuli. To verify the effect of multimodal integration, we adjusted the sound decibel level (by controlling PWM duty cycle) for clicks and the brightness (by controlling PWM amplitude) for flashes to achieve similar performance levels under unimodal conditions. We then examined whether there was an enhancement effect under multimodal stimulation with these parameter settings.

#### Confidence-proved task

In Fig. 3E, mice initiated each trial by licking the central spout, while suppressing licking of the lateral spouts for a brief period. Following this, a sound stimulus ranging from 8 kHz to 32 kHz was emitted from the top buzzer for a duration of up to 1000ms. During the first 250ms of the stimulus, the mice had to inhibit any early licks. Failure to do so triggered 100ms noise cue, prematurely terminated the current trial, and imposed a timeout penalty lasting 3 to 7 seconds. The mice’s first choice of either the left or right spout after the 250ms sample period was recorded as their decision, ending the stimulus. If the choice was correct, a small reward (approximately 0.25μL) was delivered at the corresponding spout after a brief delay. The mice then had to confirm their decision by licking the rewarded spout within 1000ms to receive a second reward (0.25μL); failure to confirm was treated as a no-response trial. During the delay, mice were required to maintain their choice, refraining from licking any other spout. Licking a non-chosen spout during the delay was interpreted as forfeiture of the decision, resulting in the termination of the trial. If the central spout was licked during this period, the trial ended immediately and a new trial began. For correct choices, the delay duration was sampled from an exponential distribution with a maximum of 5000ms, a minimum of 500ms, and an average of 1000ms. For incorrect choices, the delay was fixed at 20 seconds, with no additional error feedback provided; mice had to lick the other spouts twice to terminate the trial. Trials where mice did not actively terminate within 20 seconds were classified as no-response trials. During the sample period, the task was to discriminate whether the sound frequency exceeded 16 kHz, indicating a right spout choice, or was below 16 kHz, indicating a left spout choice. The frequency range was logarithmically partitioned around the 16 kHz midpoint into nine intervals: {-1, –0.6, –0.2, –0.1, 0, 0.1, 0.2, 0.6, 1}. Negative values corresponded to a left choice, positive values to a right choice, and 0 represented a random choice between left and right. During the testing phase, a certain percentage of the trials were designated as probe trials: when the mice made a correct choice, the delay was set to 20 seconds, and the first reward was omitted. The mice were required to lick another spout to terminate these trials.

#### Continual training

In Fig. 4A, mice initiated a new trial by licking the central spout, followed by a brief delay before entering the sample period. During the first 350ms of both the delay and sample period, the mice were required to refrain from licking the lateral spouts; doing so resulted in a noise cue and a timeout penalty. After the first 350ms of the sample period, the first lick on either lateral spout was recorded as the mouse’s choice for that trial, marking the end of the sample period. A correct choice resulted in 0.25μL water reward at the corresponding spout, while an incorrect choice triggered a noise cue and a timeout penalty. The stimuli presented during the sample period followed a specific sequence: sound frequency discrimination (3 kHz vs. 12 kHz), followed by reversed sound frequencies (12 kHz vs. 3 kHz), sound direction (left vs. right), reversed sound direction (right vs. left), and finally, light direction (left vs. right). Progression to the next stimulus condition occurred only when the mice achieved and maintained a performance accuracy above 75% in the previous discrimination task.

#### DMS

In Fig. 4B, mice initiated a trial block by licking either the left or right spout. After a brief delay, they entered a 500ms sample period, during which a sound stimulus of either 3 kHz or 12 kHz was emitted from the top buzzer. Following this, a 1-second delay period began, during which licking either the left or right spout caused the trial to pause for 300ms. The delay period was followed by a 500ms test period, with another 3 kHz or 12 kHz sound stimulus delivered from the top buzzer. This was followed by a second 1-second delay period, during which the mice were again required to refrain from licking the lateral spouts. A brief response cue then indicated that the mice could express their choice by licking either the left or right spout. A correct choice resulted in 0.25μL water reward, while an incorrect choice led to a noise cue and a timeout penalty. When the sound frequencies presented during the sample and test periods were identical, the correct choice was the left spout. When the frequencies differed, the correct choice was the right spout. During the testing phase, the durations of the two delay periods were independently and randomly sampled from {1, 1.5, 2, 2.5, 3s}.

#### d3AFC

In Fig. 4C, a paradigm similar to that described in Fig. 2A was used, but with three possible choices. Mice could initiate a trial block by licking any of the three spouts. The task required them to discriminate between three different sound frequencies: 8 kHz, 16 kHz, and 32 kHz, with the corresponding spouts being left, center, and right, respectively, to receive a reward. During the testing phase, stimuli were logarithmically spaced around the 16 kHz midpoint, with values expanded to {-1, –0.75, –0.5, –0.25, 0, 0.25, 0.5, 0.75, 1}.

Frequencies corresponding to {-1, –0.75, –0.5} indicated a left spout choice; {-0.25, 0, 0.25} indicated a center spout choice; and {0.5, 0.75, 1} indicated a right spout choice. In **Fig. S5C**, for some mice that did not undergo the psychometric curve testing phase, the left and right stimulus-choice associations were reversed after they had learned the initial task, while the center stimulus-choice association remained unchanged. Whenever the mice achieved a 75% accuracy rate across all three trial types and maintained it for some time, the left-right stimulus-choice associations were immediately reversed again. Mice could only detect these reversals through trial feedback, with no additional cues indicating changes in task rules.

#### Delayed Context-dependent task

In Fig. 4D, mice triggered a new trial by licking the central spout. After a brief delay, 500ms rule-indicating clicks stimulus was emitted from the top buzzer. This was followed by a delay period during which the mice were required to suppress licking the lateral spouts; otherwise, the current trial would end, accompanied by a noise cue and timeout penalty. Subsequently, 200ms auditory cue (3 kHz or 12 kHz) was delivered from either the left or right buzzer. Mice could report their choice 100ms into this cue. A correct choice resulted in 0.25μL water reward at the chosen spout, while an incorrect choice led to a noise cue and timeout penalty.

The click stimulus represented different task rules based on its frequency: 16 Hz clicks indicated that the mice should make their choice based on the direction of the sound, while 64 Hz clicks indicated that the reward was associated with the sound frequency. When the trial required attention to sound direction, the mice needed to choose based on the sound source location (left buzzer for left spout, right buzzer for right spout), ignoring the sound frequency. Conversely, for trials requiring attention to sound frequency, the mice had to discriminate the sound frequency (3 kHz for left, 12 kHz for right) to make their choice. The context and sample periods were separated by a 250ms delay period. Trials where the correct choices based on direction and frequency aligned were termed “coherent trials,” while those where they conflicted were termed “conflict trials.” Trials involving only one modality of stimulus were termed “unimodal trials” (for single sound direction trials, the frequency was set at 6 kHz; for single frequency trials, the sound was delivered from the top buzzer). Trials were randomly presented as 10% unimodal, 70% conflict, and 20% coherent, with mice required to achieve 75% accuracy on conflict trials.

During the testing phase, the delay duration was randomly chosen from a range of 200ms to 2000ms. Additionally, the click rate, representing the context, was extended to {16, 20, 25, 32, 40, 50, 64 Hz}, where rates below 32 Hz indicated attention to sound direction and rates above 32 Hz indicated attention to sound frequency. For trials with a 32 Hz click rate, the reward choice was randomly linked to either sound direction or frequency.

**Supplementary Fig. S1.**
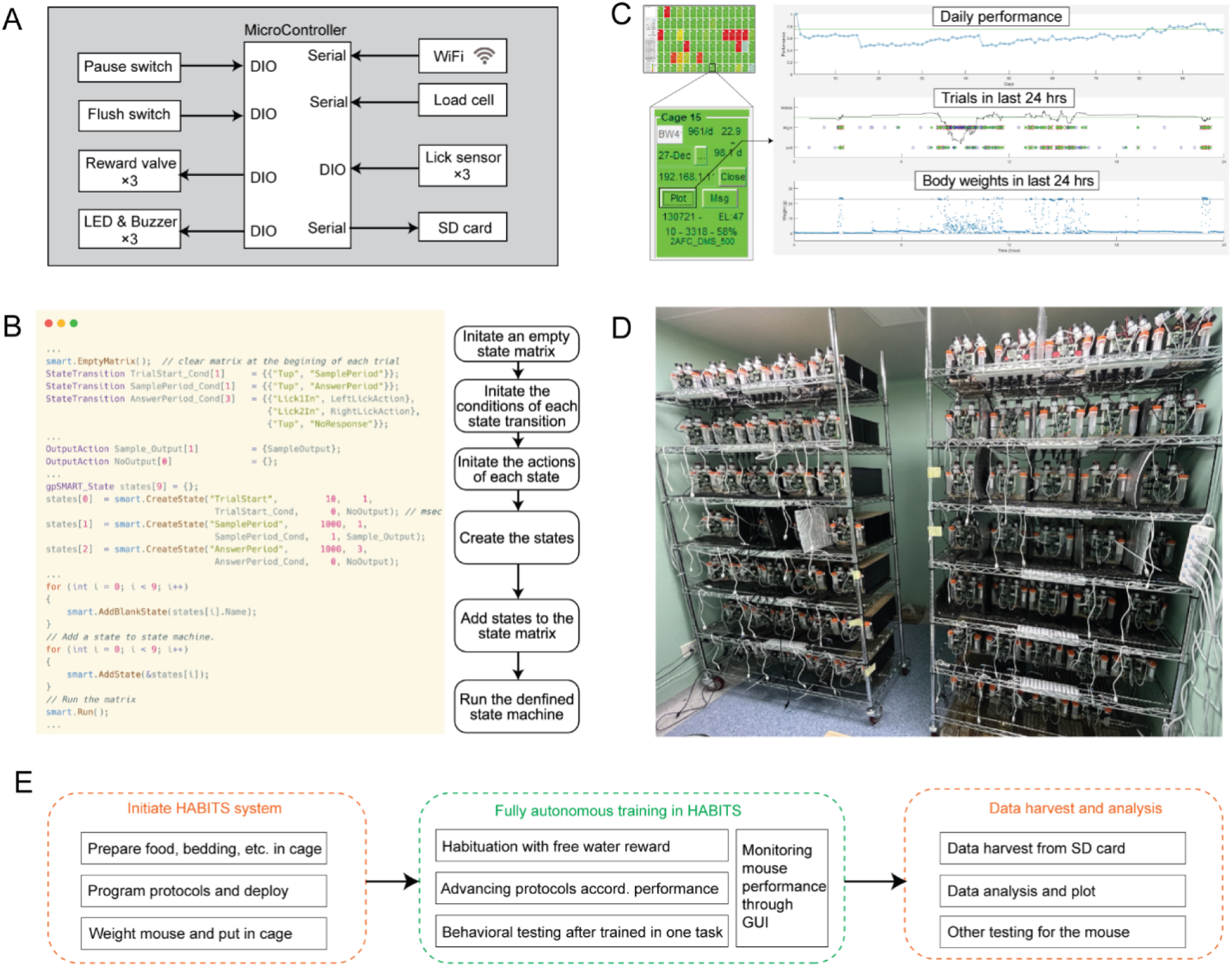
HABITS system. (**A**) Block diagram of control system of HABITS, showing peripherals connected with microcontroller through digital input/output (DIO) or serial port. (**B**) Graphic user interface (GUI) of a specific cage (left, magnified) and data plot window (right) when click ‘plot’ button in the GUI, showing daily performance in all previous days, trial performance (green for correct and red for error trials) in last 24 hours, and body weight data in last 24 hours. (**C**) Example protocol programs for HABITS. (**D**) Around 100 HABITS are packed on standard racks for large-scale mouse behavioral testing. (**E**) Workflow pipeline for HABITS, showing fully autonomous mouse behavioral training after initialization of HABITS, before data harvest from SD card for analysis.

**Supplementary Fig. S2.**
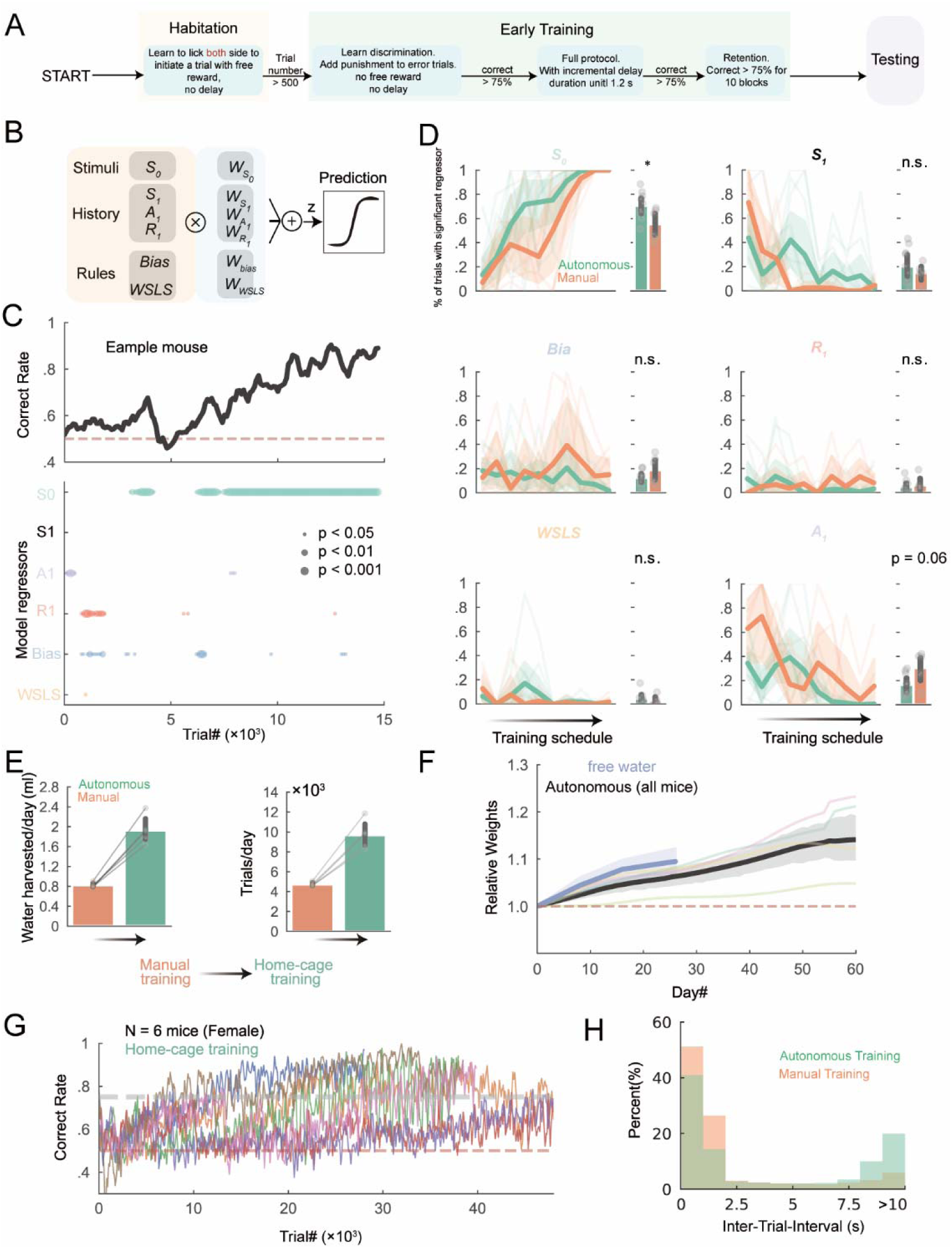
Autonomous versus manual training in home-cage. (A), Flow chart of the task training protocol in home-cage (Materials and Methods). (B), Logistic regression model. (C), top, behavioral performance of example mouse in the autonomous training. Bottom, the significance of individual regressors; Circle size corresponds to p values; The significance of a regressor is evaluated by comparing the prediction of the full model to a partial model with the regressor of interest excluded. *p*-values are based on cross-validation t-test (Materials and methods). (D), percentage of trials significantly predicted by different regressors during task learning. Cycles and light lines, individual mice; Bars and bold lines, average across mice; Shades and error bars, 0.95 CI. *, p<0.05, n.s., p>0.05, two-sided Wilcoxon rank-sum tests. (E), averaged water harvested per day (left) and number of trials per day (right) changing from manual to autonomous training in home-cage. Cycles, individual mice; Bar plot and error bar, mean and 0.95 CI across mice. (F), averaged relative body weights as a function of training days for free water (blue) and all d2AFC training mice (black). Shaded area shows 95% CI. (G), performance of all 6 female mice performing d2AFC task in home-cage automatically. (H), The histogram of inter-trial-interval for both autonomous and manual training in HABITS.

**Supplementary Fig. S3.**
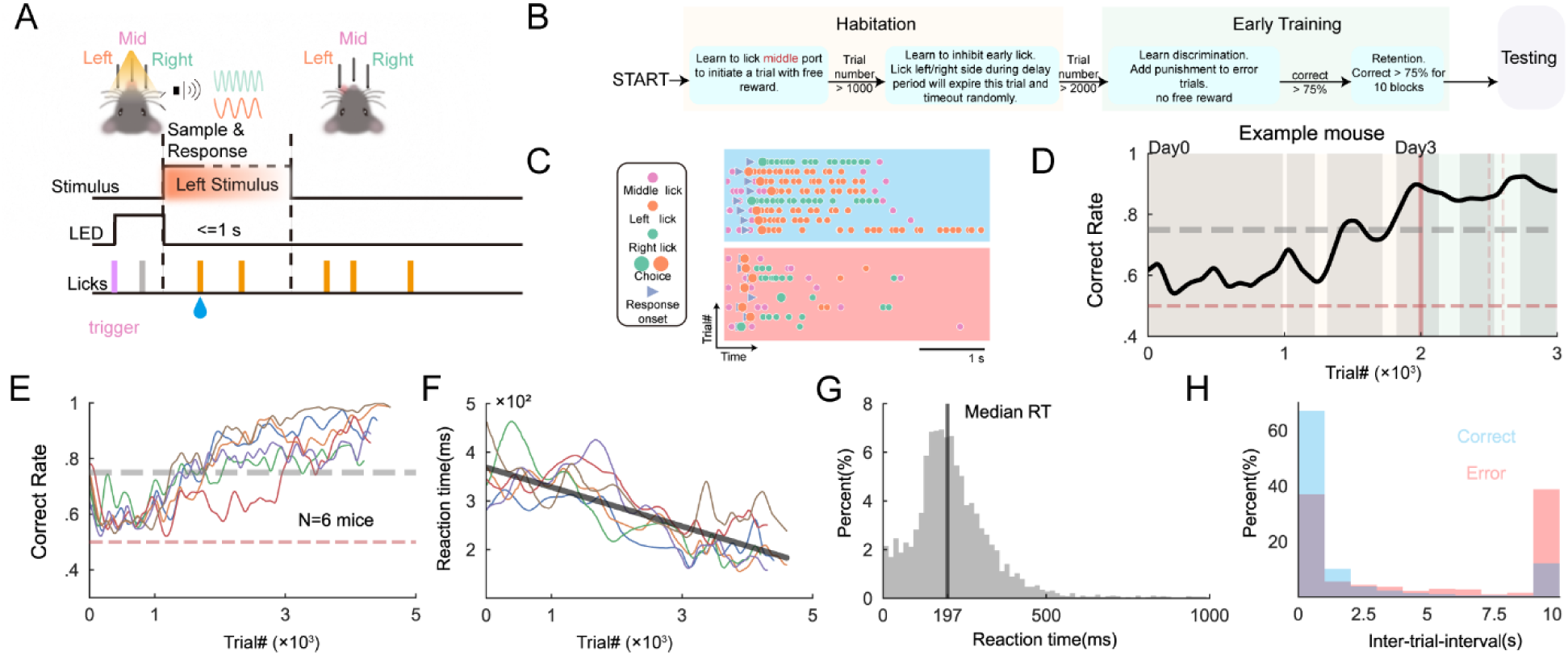
Reaction time based 2AFC task training in home-cage automatically. (**A**), task structure of RT-based 2AFC task. (B), Flow chart of training protocol in home-cage. (C), conditioned behavioral data of example trials for correct (blue block) and error (red block) choice. (D), performance of example mouse performing task in home-cage. The color of background corresponding to (B). Grey blocks indicate dark cycle. Grey dash line, the criterion performance. Red horizontal dash line, chance performance level. (E), correct rate of all mice. (F), reaction time of all mice. Black line fitting to all mice from the onset to the end of training. (G), histogram of reaction time. Data collected from all mice. The bold vertical line represents the median of RT. (H), conditioned histogram of inter-trial-interval (ITI) for correct (blue) and error (red) trials.

**Supplementary Fig. S4.**
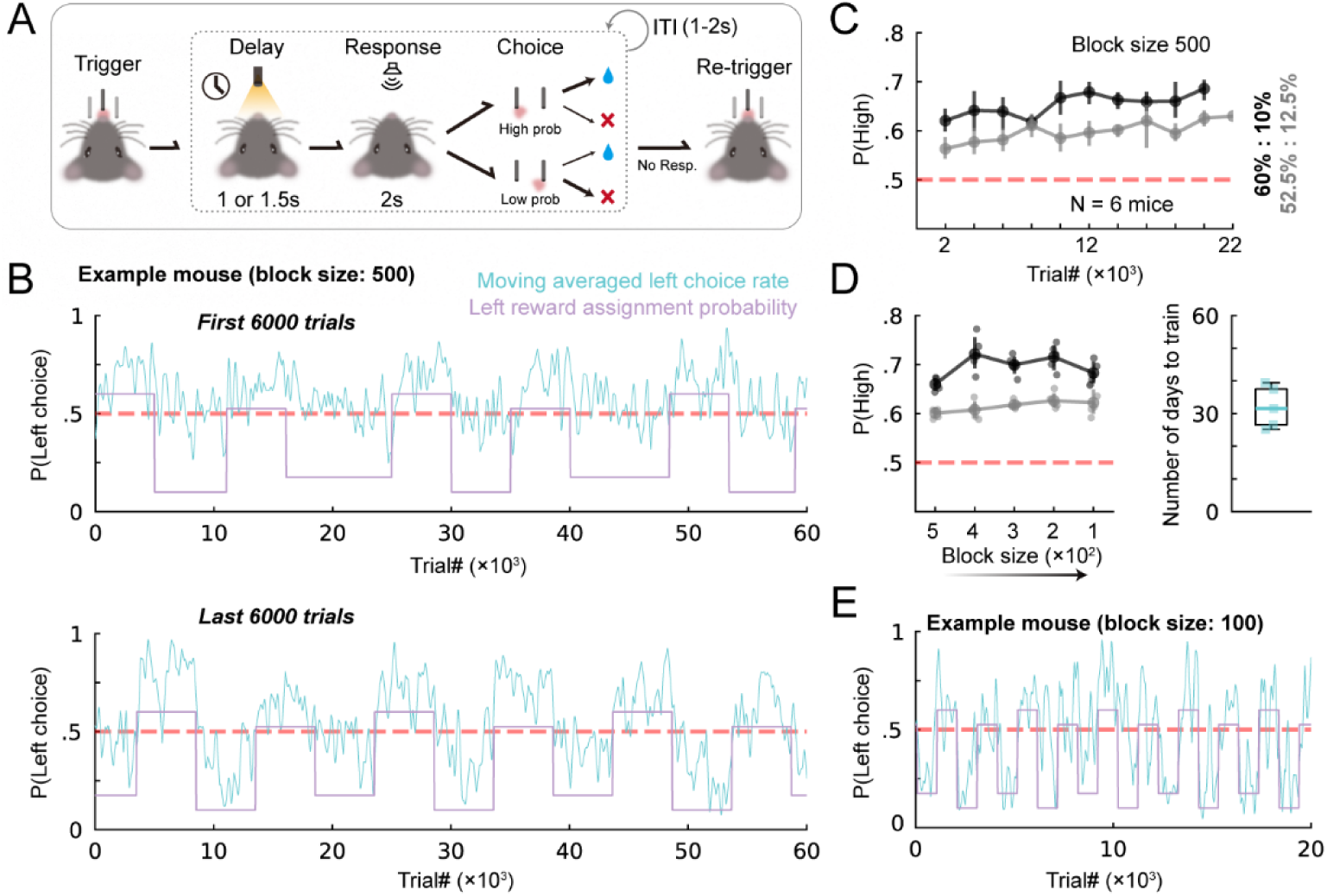
Value-based dynamic foraging task. (**A**), task structure. (**B**), example performance of a mouse in the early (top, first 6000 trials) and late (bottom, last 6000 trials) training stages with block size 500. Blue lines represent moving averaged behavioral probability of left choice within 40 trials. Purple lines show the assignment probability for left reward. (**C**), averaged probability of the choosing the lickport with the higher assignment probability (P(high)) across mice gradually increases following the number of trials. Black line indicates the assignment probability for left and right lickports is 60% (grey line, 52.5%) and 10% (grey line, 17.5%), respectively. Dots and errorbar, mean and 95% CI. (**D**), left, averaged P(high) across mice follows training sub-protocols with different block size. Right, the number of days to complete all training protocols from block size 500 to 100. Square dots indicate individual mice. (**E**), same as (B) but data collected from the sub-protocol with block size 100.

**Supplementary Fig. S5.**
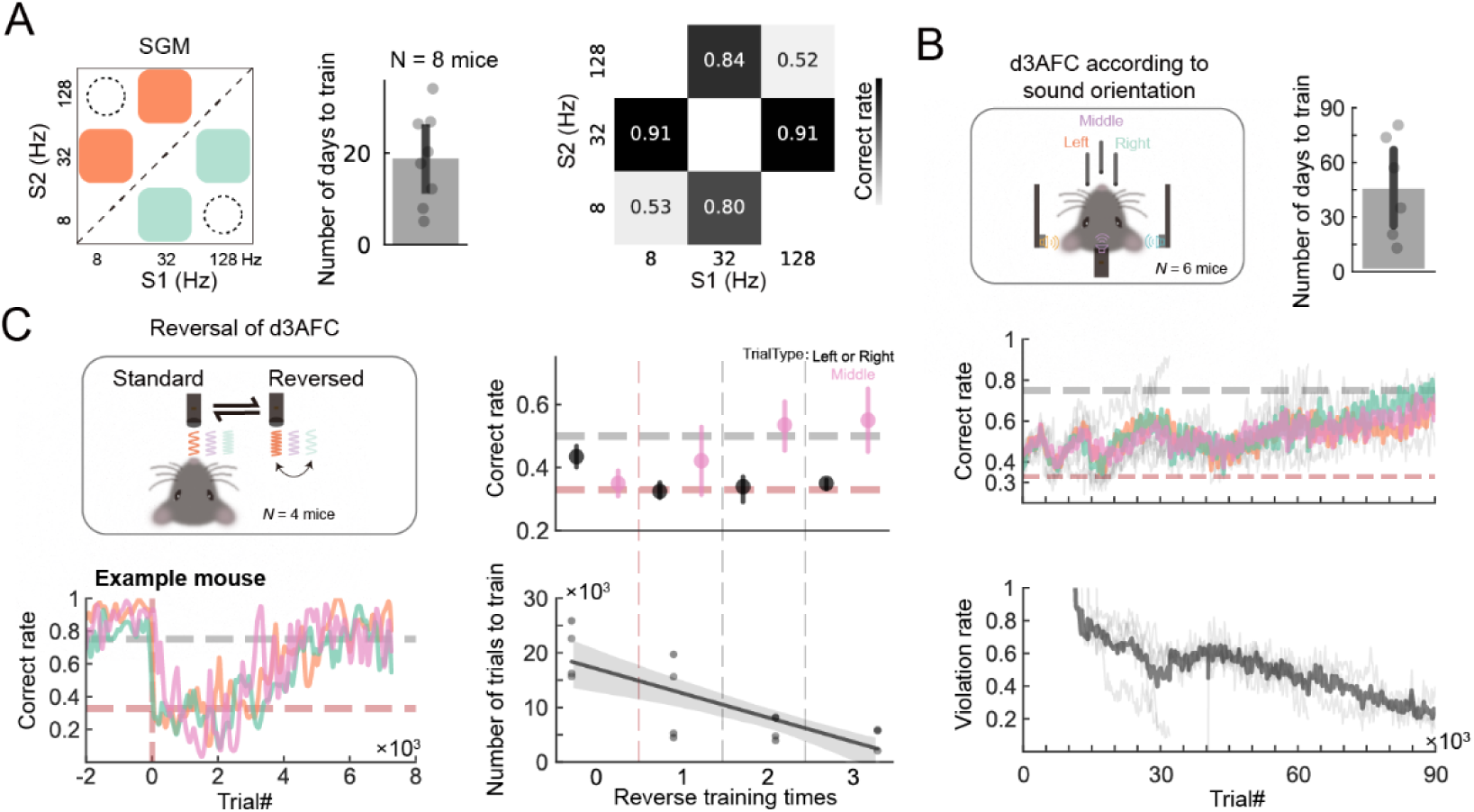
Other complex cognitive behavioral tasks training in home-cage automatically. (A), Left, stimulus generation matrix of working memory task. Middle, number of days to train. Right, correct rate for SGM. Values lying in the diagonal line corresponding to the correct rate of probe trials. (B), Top, d3AFC task according to sound orientation and number of days to reach the criterion performance. Dots indicate individual mice. Performance (Middle) and earlylick rate (bottom) of all mice performing the d3AFC task. Red dash line, chance performance level; Grey dash line, the criterion performance. (C), Left, contingency reversal of d3AFC task according sound frequency (top) and performance of an example mouse (bottom). Right, averaged correct rate across all mice for different reverse times (top). Number of trials needed to learn as a function of reverse training times (bottom). Dots, individual mice. Line and shades, linear regression.

**Supplementary Fig. S6.**
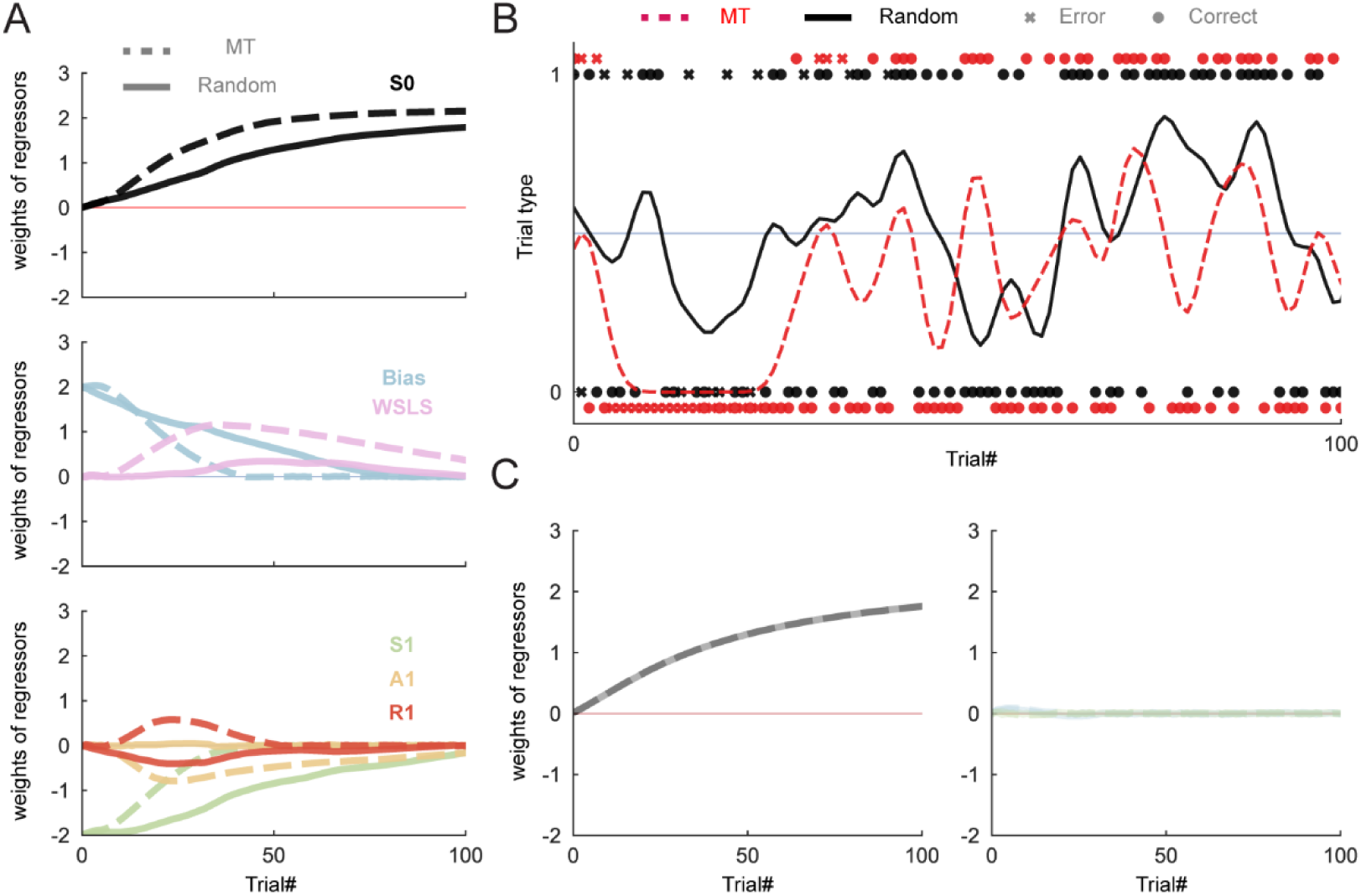
Simulation of machine teaching algorithm in decision-making scenario. (**A**). the weight of regressors in an ideal learner vary during learning a 2AFC task. Note that the initial weights of bias and S1 regressors are not zero. (**B**), the presented trial types generated by random (black) and MT (red) during entire training process. (**C**), same as (A) but weights of all regressors begin at zero.

**Supplementary Fig. S7.**
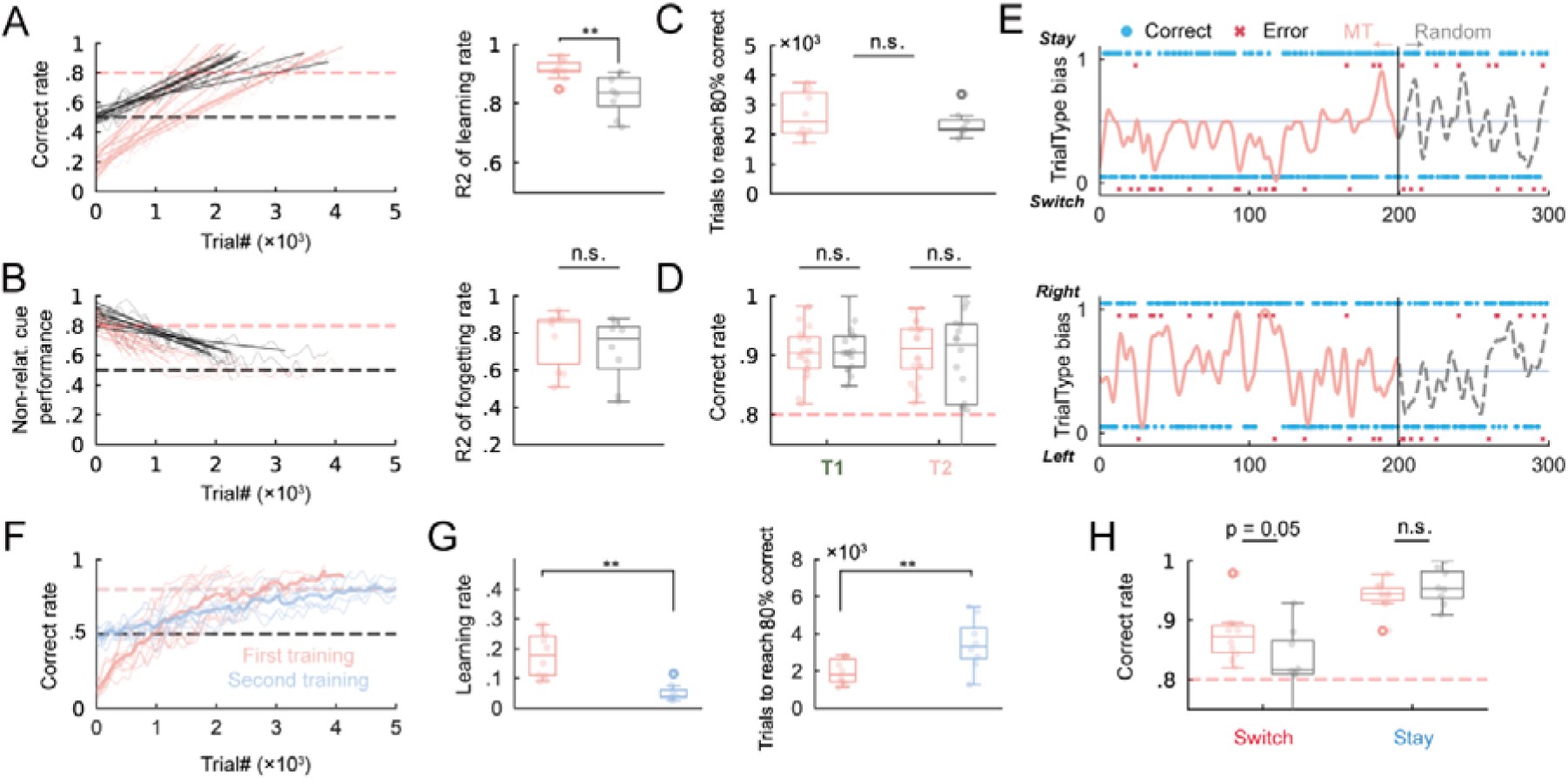
Details of behavioral analysis for multi-dimensional tasks. (**A**), left, linear regression between trial number and correct rate in task requiring mice attend to sound frequency. Right, the R-square of every individual linear regression. (**B**), same as (A) but for performance following non-relative cue. (**C**), the number of trials to reach criterion performance for MT and random group. (**D**), performance of both grouped mice in T1 and T2 protocol. n.s., no significant. two-sided Wilcoxon rank-sum tests. (**E**), the presented individual trials with Stay/Switch (top) and Left/Right (bottom) trial type generated by MT (L3) and Random (T2). (**F**, **G**), After mice were trained by MT as in fig. 6A, they were intermediately set the training protocol to the beginning and retrained with randomly generated trial sequence. We compared correct rate of trials with sound frequency stimulus in the first and the second training, presented in (F). (G) shows the learning rate (left) and training efficiency (right) of the first and the second training processes. **, p<0.01; two-sided Wilcoxon signed-rank tests. **(H**), correct rate of both grouped mice for stay and switch trials in T2 protocol. n.s., no significant. two-sided Wilcoxon rank-sum tests.

**Supplementary Table S1.** Building materials of HABITS.

| **Parts** | **Quantity** | **Model and Supplier** | **Price\*** |
| --- | --- | --- | --- |
| Acrylic board | N/A | Custom | \$ 8.47 |
| 3D printing | N/A | Sogaworks.com | \$ 9.88 |
| Microcontroller | 1 | Teensy 3.6, PJRC Inc. | \$ 36.43 |
| SD card | 1 | SDQUNC-16GB, SanDisk Inc. | \$ 3.94 |
| Load cell and driver module | 1 | YZC191 and HX711,<br>Guangce.com | \$ 3.57 |
| Licking sensor module | 1 | MPR121, Taobao.com | \$ 0.49 |
| Wi-Fi module | 1 | ESP8266, Espressif Inc. | \$ 0.76 |
| LED | 3 | GS5050RGB and<br>GS3528UWC, Guangsheng<br>Inc. | \$ 0.14 |
| Buzzer | 3 | PS1740P02, TDK Inc. | \$ 1.27 |
| Solid State Relay | 2 | SDD-5HB, XYF Inc. | \$ 9.32 |
| Peristaltic Pump | 2 | KPP-DCL-S01W, kamoer Inc. | \$ 20.0 |
| Power Adapter | 1 | ABLK-T1201, Aobaolike Inc. | \$ 1.40 |
| Others | N/A | N/A | \$ 3.07 |
| **Total** | | | **\$ 98.74** |

\*Price was originally in Chinese yuan and converted to US dollar with exchange rate of 7:1.

**Supplementary Movie S1.** Free moving mouse performing task in HABITS.

**Supplementary Movie S2.** The 24-hour activities of mouse living in HABITS.

